# Social information drives ecological outcomes among competing species

**DOI:** 10.1101/604595

**Authors:** M.A. Gil, M.L. Baskett, S.J. Schreiber

## Abstract

Through its behavior, an organism intentionally or unintentionally produces information. Use of this ‘social information’ by surrounding conspecifics or heterospecifics is a ubiquitous phenomenon that can drive strong correlations in fitness-associated behaviors, such as predator avoidance, enhancing survival within and among competing species. By eliciting indirect positive interactions between competing individuals or species, social information might alter overall competitive outcomes. To test this potential, we present new theory that quantifies the effect of social information, modeled as predator avoidance signals/cues, on the outcomes from intraspecific and interspecific competition. Our analytical and numerical results reveal that social information can rescue populations from extinction and can shift the long-term outcome of competitive interactions from mutual exclusion to coexistence, or vice versa, depending on the relative strengths of intraspecific and interspecific social information and competition. Our findings highlight the importance of social information in determining ecological outcomes.

## Introduction

The mere presence and even simple behaviors of an individual animal produce sensory information that becomes publicly available to surrounding individuals (Danchin et al. 2004, Dall et al. 2005, Goodale et al. 2010). Such ‘social information’ has long been a central topic of interest in select study systems in which individuals intentionally produce signals (Templeton and Giraldeau 1995, Magrath et al. 2015). However, recent empirical and theoretical evidence from various systems indicates that social information use extends far beyond intentional signaling and appears to be a general phenomenon in systems in which individuals that cohabit a landscape share needs (Seppänen et al. 2007, Goodale et al. 2010, Gil et al. 2017, Gil and Hein 2017, Kane and Kendall 2017, Gil et al. 2018). Perhaps the best studied and most common individual need that is enhanced by social information is predator avoidance: alarm calls warn of approaching predators in avian and primate systems (Zuberbühler 2001, Danchin et al. 2004, Magrath et al. 2015), postures, evasive movements, or the use of predator-free space inadvertently provide information on the proximity of threats in avian, mammalian, and fish systems (Griffin 2004, Schmitt et al. 2016, Gil and Hein 2017), and even plants can use chemical cues from damaged neighboring plants to induce defenses to protect against herbivores (Karban et al. 2000, Dicke and Bruin 2001). Because social information typically enhances the fitness of receiving individuals, and, because any individual in a population can repeatedly receive such benefits, social information could affect the dynamics of populations (Gil et al. 2018). Thus, understanding the degree to which social information can affect population dynamics is a pressing question in ecology.

Social information creates the potential for indirect positive interactions within and across species and might drive positive density dependence. Positive density dependence (i.e., an ‘Allee effect’) occurs when a greater density of individuals in a population enhances the growth rate of that population (Courchamp et al. 1999, Stephens et al. 1999). This simple process can drive profound changes to the dynamics of a population, affecting not only a population’s carrying capacity but also its likelihood of sudden change or collapse (Stephens and Sutherland 1999, Schreiber 2003). For example, under positive density dependence, loss of individuals (e.g., due to harvesting) can become increasingly detrimental to a population, even leading to negative population growth when a population falls below a critical threshold (i.e., a ‘strong Allee effect’; Stephens et al. 1999). Positive density dependence and the critical population thresholds they can cause are putatively common though difficult to rigorously identify in natural systems and are, therefore, of particular interest to natural resource conservation and management (Stephens and Sutherland 1999, Berec et al. 2007, Gregory et al. 2010). Positive density dependence has been classically attributed to non-information-mediated mechanisms, such as mate limitation or habitat amelioration, and to information-mediated mechanisms in species that form cohesive groups, such as flocks or schools (Courchamp et al. 1999, Stephens and Sutherland 1999, Stephens et al. 1999, Gascoigne and Lipcius 2004b). Yet, positive density dependence can arise due to social information regardless of whether or not individuals form cohesive groups or are conspecifics. Social information typically enhances individual survival or reproduction and increases with the density of information-producing individuals (Kenward 1978, Jackson et al. 2008, Kazahari and Agetsuma 2010, Lister 2014, Berdahl et al. 2016, Gil et al. 2017, Gil et al. 2018).

Social information use is most likely between individuals in similar guilds (e.g., those on the same trophic level with shared predators) and, thus, typically occurs in the context of intraspecific and interspecific competition for resources. As a negative interaction, competition counters the effects of social information. Effects of both competition and social information are density dependent, but in opposing ways. Social information typically is most beneficial at low to intermediate population densities, where information is less redundant or its benefits less ephemeral. In contrast, competition typically is most detrimental at higher densities, where resources are more limited (Gil et al. 2018). Thus, we expect social information to have stronger net per capita effects when population densities are low, as they often are in human-altered landscapes (Courchamp et al. 1999). Nonetheless, to measure the net impact of social information requires knowing the strength of competition. Furthermore, competition and the exchange of social information can occur to varying degrees both within species and across species (Monkkonen et al. 1999, Seppänen et al. 2007, Goodale et al. 2010). Therefore, to understand the ecological consequences of social information requires that we examine the joint effects of intraspecific and interspecific social information and intraspecific and interspecific competition.

Population models offer a framework through which to explore the population- and community-level consequences of social information use in wild animals. Classic models that measure the demographic effects of predator functional response, positive density dependence, and facilitation provide conceptual precursors to the study of social information. Noy-Meir (1975) showed that the deceleration of a generalist predator’s attack rate across low prey densities (a Type II functional response (Holling 1966) can generate a strong Allee effect (Noy-Meir 1975)). In this model, and in most population models, this deceleration of the predator’s attack rate with prey density is attributed to properties of the predator (e.g., satiation, handling time; (Oaten and Murdoch 1975)). However, this deceleration could be driven by properties of the prey themselves (i.e., if more prey better help one another avoid predation). More recently, models exploring the demographic effects of mutualism have shown that even when positive interspecific interactions are constrained to low and narrow population density ranges, they can quantitatively and qualitatively affect the fate of one or both interacting populations (Hernandez 1998, Hernandez and Barradas 2003, Zhang 2003, Zhang et al. 2007, Hernandez 2008, Holland and DeAngelis 2009, Holland and DeAngelis 2010). Social information provides a possible mechanism for density dependent mutualism, but with the added complexity of being shared not only between species but also within species (Gil et al. 2018). The two existing population models that explicitly account for social information (Schmidt et al. 2015, Schmidt 2017) focus on the case of enhanced breeding habitat selection among conspecifics and show that social information can drive strong Allee effects. Evaluating whether such critical thresholds occur in multi-species systems with social information requires building on this theory to explicitly model social information in a multispecies context.

Here, we use models of a single species and of competing species to build a theory of the demographic consequences of social information use in wild animals. We focus on the widespread use of social information about predators. We modify a framework developed in Gil et al (2018), where we demonstrated that social information can alter qualitative expectations for population and community dynamics in specific cases, to thoroughly and comprehensively quantify, and develop metrics for, when such qualitative changes are expected to occur. We first quantify the intraspecific effects of this social information using a reparameterization of the classic Noy-Meir model to address the question: under what conditions does this common form of social information affect the existence of critical thresholds, equilibrium densities or persistence of a population? We then expand to a two-species population model, in which competition and the exchange of social information can occur within and between species, to address the question: how does social information affect the nature and outcome of species interactions? Our study reveals that social information can alter competitive outcomes and generate multiple alternative stable states in a predictable manner depending on the relative strengths of intra- and interspecific competition, intra- and interspecific social information, and predation. Our modeling framework is general, meaning it is not system-specific, and our findings lay the groundwork for further theoretical and empirical investigation of how social information scales up to affect the ecology and conservation of natural systems.

## Methods

### Effects of social information on single species dynamics

To lay the groundwork for our two-species model, we begin with the dynamics of a single species with population size *N* that exhibits logistic growth, determined by intrinsic per-capita growth rate *r* and intraspecific competition coefficient *α* (the carrying capacity is 1/α). The population also experiences mortality due to a generalist predator at a maximal rate *p*, and per-capita mortality decays with prey density, through the sharing of social information (e.g., alarm calls, evasive movements; Danchin et al. 2004, Goodale et al. 2010, Magrath et al. 2015), and the per-capita strength of social information *b*. In other words, social information reduces per-capita predation rates by extending capture time. Thus, the single species dynamics are

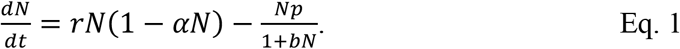

Here, per-capita predation risk saturates at high prey densities because prey reach the maximal per-capita benefit of social information on predator avoidance (i.e., diminishing returns on information due to redundancy, ephemeral benefits or occlusion of information motivate a saturating functional form; Kenward 1978, Seppänen et al. 2007, Jackson et al. 2008, Lister 2014, Berdahl et al. 2016). The model assumes the predator population size remains constant, independent of prey density, *N*, and that the predator has a linear functional response. However, if we assume the predator exhibits a Type II functional response, then we still get the same functional form of the predation term in Eq. 1 (Type II) but with new parameters (see Appendix S1 for details). Furthermore, while Eq. 1 allows social information to drive the predation rate to zero, an unlikely outcome in most natural systems, this equation is mathematically equivalent to a functional form in which social information causes predation rate to level off at a nonzero value, determined by an additional parameter (see Appendix S1 for details). Thus, all of our results about Eq. 1 also apply to models with a Type II predator functional response and a minimal predation level even when social information is high.

### Effects of social information on competing species with shared predators

We expand upon the model presented in Eq. 1 to measure how social information can affect the long-term dynamics of two competing species. We follow population sizes *N_i_* of each species *i*, where within-species population growth, *r*_i_, density dependence, *α*_ii_, and a maximal per-capita predation rate, *p_i_*, follow the same dynamics and notation as Eq. 1, but these species compete with one another at a rate *α_ij_*, which represents the per-capita negative effect of the *j*-th species on the i-th species, where *i* ≠ *j*. Both species experience per-capita mortality due to predation at a rate that decays with increasing densities of both species (the two-species analog of the mortality term in Eq. 1); i.e., both conspecifics and heterospecifics share and use social information (e.g., from alarm calls or evasive movements) to enhance predator avoidance (Danchin et al. 2004, Goodale et al. 2010, Magrath et al. 2015). Note that this functional form of mortality could also be used to model non-information-mediated interactions, such as species-specific prey handling times by the predator, or group defenses that increase with density. Here, *b_ii_* represents the magnitude of the effect of intraspecific social information and *b_ij_* (*i* ≠ *j*) that of interspecific social information, such that the dynamics of the *i*-th species are

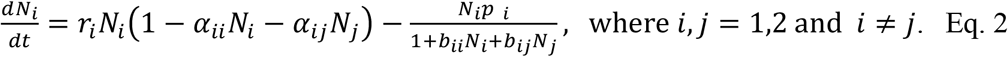

We provide a mechanistic derivation of this two-species model by considering transitions between informed and uninformed behavioral states of individuals in the community (Appendix S1). Note that Eq. 2 is equivalent to a version of the model that includes predator handling time (i.e., a Type II functional response) under special cases (Appendix S1, Eq. S3).

### Analysis of models

For the single species model in Eq. 1, we conduct a global bifurcation analysis for different values of *b*, to determine under what conditions social information about predators can alter the persistence of a population, generate a strong Allee effect, and alter the equilibrium density of a persisting population. For the mathematical and numerical analysis of the competing species model in Eq. 2, we focus primarily on the case that the species are symmetric, i.e., *r*_1_ = *r*_2_ = *r*, *α*_11_ = *α*_22_ = *α*, 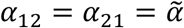, *b*_11_ = *b*_22_ = *b*, 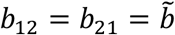, *p*_1_ = *p*_2_ = *p*. For this two-species model, we analytically derive conditions for different community outcomes and develop an analytically-based numerical method to identify all equilibria and their stability. We use these methods in conjunction with numerically computed isoclines to determine how social information affects the nature and outcome of species interactions. Specifically, we compare individual and combined effects of intraspecific and interspecific social information under different relative strengths of intraspecific and interspecific competition.

## Results

### Single-species model of social information use

In the single-species model, social information can enhance persistence likelihood, with threshold dynamics, and equilibrium population size (Fig. 1). When the intrinsic per-capita growth rate *r* is greater than the maximal per-capita predation rate *p*, the population persists at a stable equilibrium for all positive initial densities. Our mathematical analysis (see Appendix S2 for details) implies that this stable equilibrium density always increases with social information (Fig. 1b; Appendix S2: Fig. S1a).

**Fig. 1:**
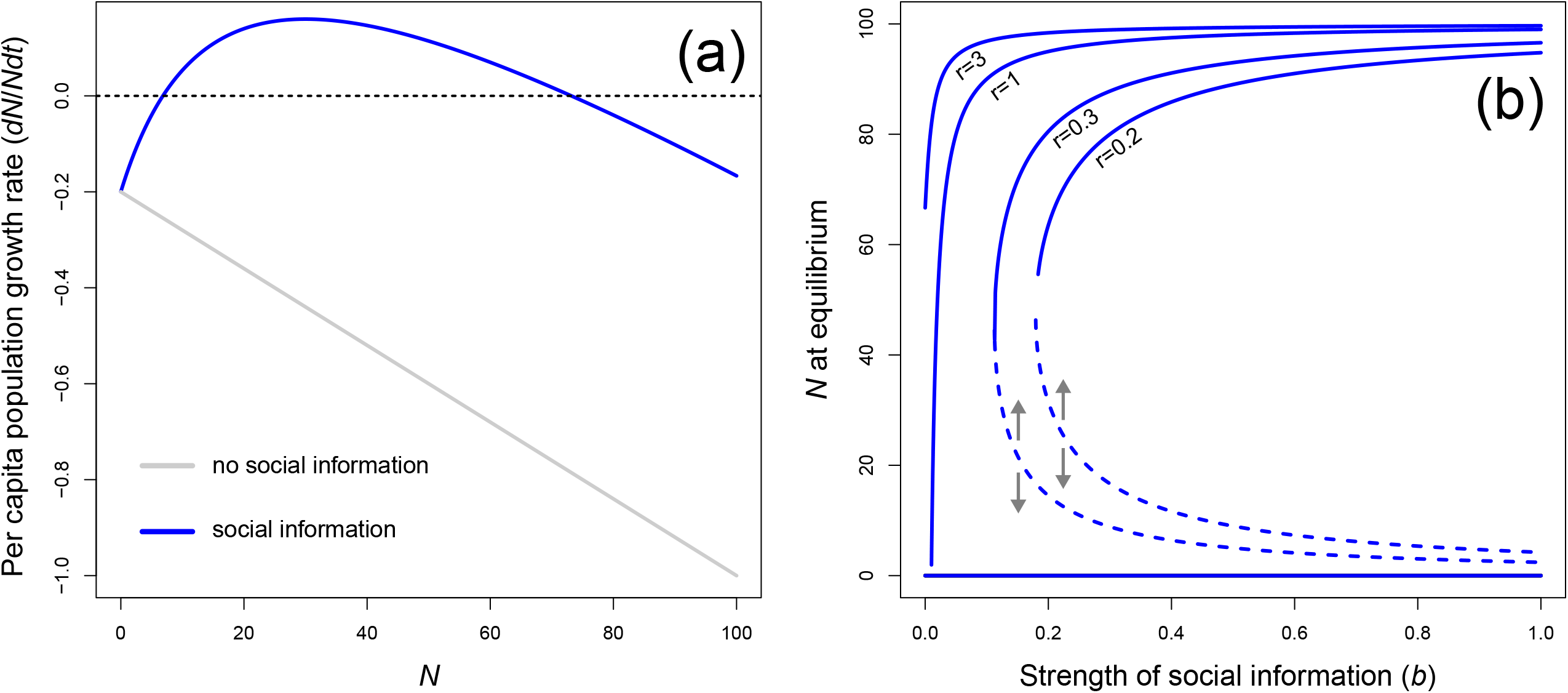
How social information can affect the dynamics of a single population. Inclusion of effects of social information on per-capita mortality due to predation can gives rise to positive density dependence in the per-capita population growth rate (a). This can expand, relative to the logistic model with predation (grey line), the conditions under which a population can persist, and it increases population size at equilibrium across a range of conditions (blue line in (a), with equilibrium population size plotted in (b)). When the predation rate exceeds the population growth rate (*p* > r), as it does in (a), social information can give rise to a strong Allee effect, causing alternative stable states (e.g., stable equilibria represented by a solid curve and a solid line at *N* = 0 for *r* = 0.2 or 0.3 in (b)). The alternative stable states are separated by unstable equilibria (e.g., the dashed curves corresponding to *r* = 0.2 and 0.3 in (b)), which represent the Allee threshold: if a population exceeds this threshold it will grow (represented by the up arrow) and if it falls below this threshold it will collapse (represented by the down arrow). When *r* ≥ *p*, social information simply increases the stable population size at equilibrium (shown for *r* = 1, 3 in (b)). Parameter values: *α* = 0.01, *r* = 0.8 (for a only), *b* = 0 (black line) or 0.05 (blue curve; for a only), and *p* = 1.

When the maximal per-capita predation rate exceeds the intrinsic rate of growth, the extinction equilibrium is stable, and the population tends to extinction whenever the initial population density is low. Social information, however, can generate a strong Allee effect and allow the population to persist whenever the maximal per-capita predation rate lies below threshold value (see Appendix S2 for details)

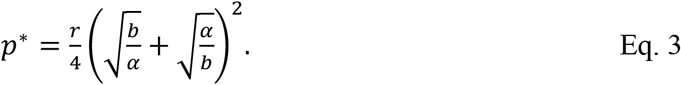

Equation 4 implies that social information that is strong relative to competition (*b* > *α*) can prevent extinction for a population at sufficiently high density (Fig. 1b; Appendix S2: Fig. S1b). When the maximal per-capita predation rate, *p*, exceeds the critical threshold *p*\*, the population goes extinct for all initial population densities (Appendix S2: Fig. S1c); population persistence is not possible. When there is a strong Allee effect (i.e. *r* < *p* < *p*\*), social information has opposing effects on the unstable equilibrium (below which the population tends to extinction) and the positive stable equilibrium. The population density at the unstable equilibrium decreases with increasing social information, while the density at the stable equilibrium increases (see Appendix S2 for a proof; compare dashed and solid curves in Fig.1b). Thus, with more social information, a population can recover from larger disturbances that reduce their densities and can ultimately approach higher densities. This pattern of social information causing positive density dependence and rescuing populations under high predation is robust to the functional form of the reduction of predation due to social information (Appendices S1 & S2; Appendix S2: Fig. S1 & S2, including the functional form used in Gil et al 2018, where the results here indicate the level of social information necessary to produce the type of qualitatively distinct behavior in the example case study of Gil et al. 2018 Box 2).

### Two-species model of social information use

Whether and how social information changes the qualitative outcome from competition within and between species depends on its strength and type. Our mathematical analyses of the two-species model (Eq. 2; Fig. 2a) when the competing species are symmetric provide information about the invasibility of the single-species equilibria and the multiplicity of equilibria on the single-species axes and on the two-species symmetric (*N*_1_ = *N*_2_) axis (see Appendix S3 for details). These analyses identify under what conditions increasing the maximal per-capita predation rate changes the ecological dynamics in two ways. First, we identify when increasing the predation rate shifts the system from coexistence via mutual invasibility (i.e. each species can invade the equilibrium determined by the other species) to mutual exclusion (i.e. each of the single species equilibria are stable), or vice versa. Second, we identify when increasing this predation rate leads to alternative stable states supporting both species or alternative states only supporting a single species. This analysis reveals that the dynamics of the system depend qualitatively on the joint effects of social information and competition via two simple net interaction indices whose form depends on the strength of social information. When both intraspecific and interspecific social information are weak (i.e., 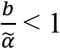 and 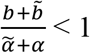; see Appendix S3 for details), the interaction index equals

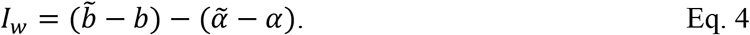

**Fig. 2:**
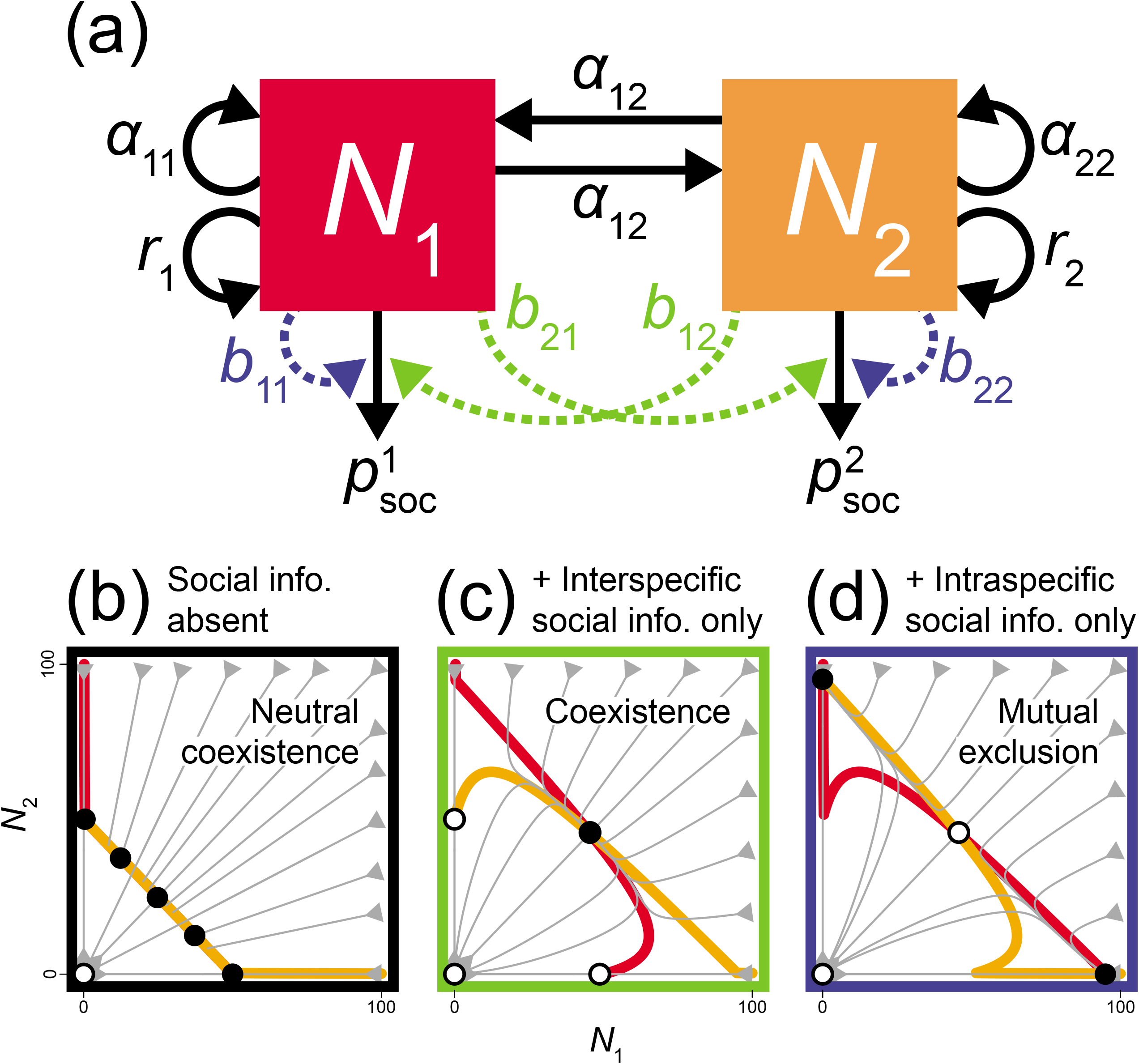
Outline of the two-species social information model (Eq. 2), including distinct effects of intraspecific social information (blue) and interspecific social information (green). (a) Boxes indicate the two population state variables, and arrows indicate dynamics and are labeled with the associated parameters. Each population exhibits logistic growth, engages in intraspecific competition (*α*_ii_) and interspecific competition (*α*_ij_), and is consumed by the same predator at a maximal per-capita rate *p_i_*. These competing species can also reduce their mortality rate due to predation by sharing intraspecific social information (*b*_ii_, in blue) and/or interspecific social information (*b*_ij_, in green); e.g., through alarm calls or evasive movements that provide early warnings of attacks). In (b)-(d), we show example phase plane plots of the competitive dynamics without social information (b), with only intraspecific social information, *I*_s_ = −1 (c), and with only interspecific social information, *I*_s_ = infinity (d). In these phase plane plots, colored lines are nullclines that indicate where each population exhibits a zero growth rate (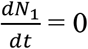: red line/curve, 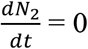: yellow line/curve; these intersect at equilibria) and grey arrows denote the trajectories populations take through time, starting from the edges of the plotted area. Open points denote unstable equilibria, and closed points denote stable equilibria. Parameter values: *r_1_* = *r_2_* = 1, *p_1_*=*p_2_* = 0.5, *α_11_* = *α_22_* = *α_12_* = *α_21_* = 0.01 with *b_11_* = *b_22_* = 0 (b, c) or 0.1 (d) and *b_12_* = *b_21_* = 0 (b, d) or 0.1 (c).

When at least one form of social information (intraspecific and/or interspecific) is strong (i.e., 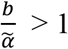 and/or 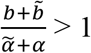; see Appendix S3 for details), the interaction index equals

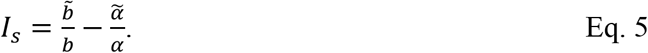

As detailed below, these net interaction indices serve two distinct purposes: 1) their signs determine the sequence of possible dynamics a symmetric system can exhibit (i.e., whether social information will push a system toward competitive exclusion or coexistence), and 2) depending on the strength of social information and predation, these indices can mark the boundary between coexistence and mutual exclusion or the boundary between persistence (of one or both species) and extinction. We first explore the effect of social information under neutral competition and then evaluate the full array of outcomes under non-neutral competition, closing with a description of the contexts in which we would expect social information to alter competitive outcomes.

### Effects of social information in competitively neutral communities

For the neutral dynamics (Fig. 2b), because 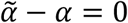 and 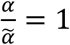, the signs of *I_w_* and *I_s_* always agree, and this index is positive only if interspecific social information is stronger than intraspecific social information (i.e., 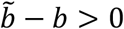, or, equivalently, 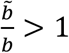). In this case, interspecific social information, which effectively decreases competitiveness between species by countering this negative effect, causes each species to have a positive per-capita growth rate when it is rare and its competitor is common (Fig. 2c). At high densities, diminishing returns of social information (e.g., due to redundancy, ephemeral benefits or occlusion of information; Kenward 1978, Seppänen et al. 2007, Jackson et al. 2008, Lister 2014, Berdahl et al. 2016) will saturate the positive effects of heterospecific density (Fig. 2c) and competition will ultimately constrain population growth. Therefore, for *I_w_ >* 0 or equivalently *I_s_* > 0, social information promotes coexistence: e.g., even weak interspecific social information (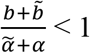; see Appendix S3 for details) shifts competitively neutral Lotka-Voltera dynamics (Fig. 2b) to coexistence (Fig. 2c). Conversely, if intraspecific social information is greater than interspecific social information (i.e., 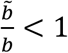), such that *I_w_* < 0 or, equivalently, *I_s_* < 0, then intraspecific social information, which effectively increases competitiveness between species by countering negative interactions within species, causes each species to have a negative per-capita growth rate when rare and its competitor is common. Therefore, even weak intraspecific social information (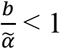; Appendix S3) shifts neutral coexistence to exclusion (Fig. 2d).

### Effects of strong social information in competitively non-neutral communities

Under non-neutral Lotka-Volterra competitive dynamics (i.e., that lead to coexistence or mutual exclusion, depending on competitive strength, given our assumption of symmetric competitors), social information interacts with the relative strengths of intraspecific and interspecific competition for symmetric species to determine the sign of *I_s_*, and the outcomes further depend on the maximal per-capita predation rate (*p*). Increasing *p* strengthens the effects of both intraspecific and interspecific social information, relative to the effects of competition and, thus, leads to different qualitative outcomes for the effect of social information on competitive dynamics. Below, we evaluate the effects of social information first when neither form (intraspecific or interspecific) is strong, then when only one form is strong, and, finally, when both forms are strong.

When neither form of social information is strong, prey populations persist only when *r* > *p*, and, in this case, the sign of *I_w_* (Eq. 4) determines whether one prey species can drive the other to extinction or the two competing species can coexist (Fig. 3a). When either form of social information is strong or both forms are strong, whether competing prey species will coexist or go extinct depends on predation level and on the sign of *I_s_* (Fig. 3b, Eq. 5), which determines the suite of possible dynamics the system can exhibit, as detailed below.

**Fig. 3:**
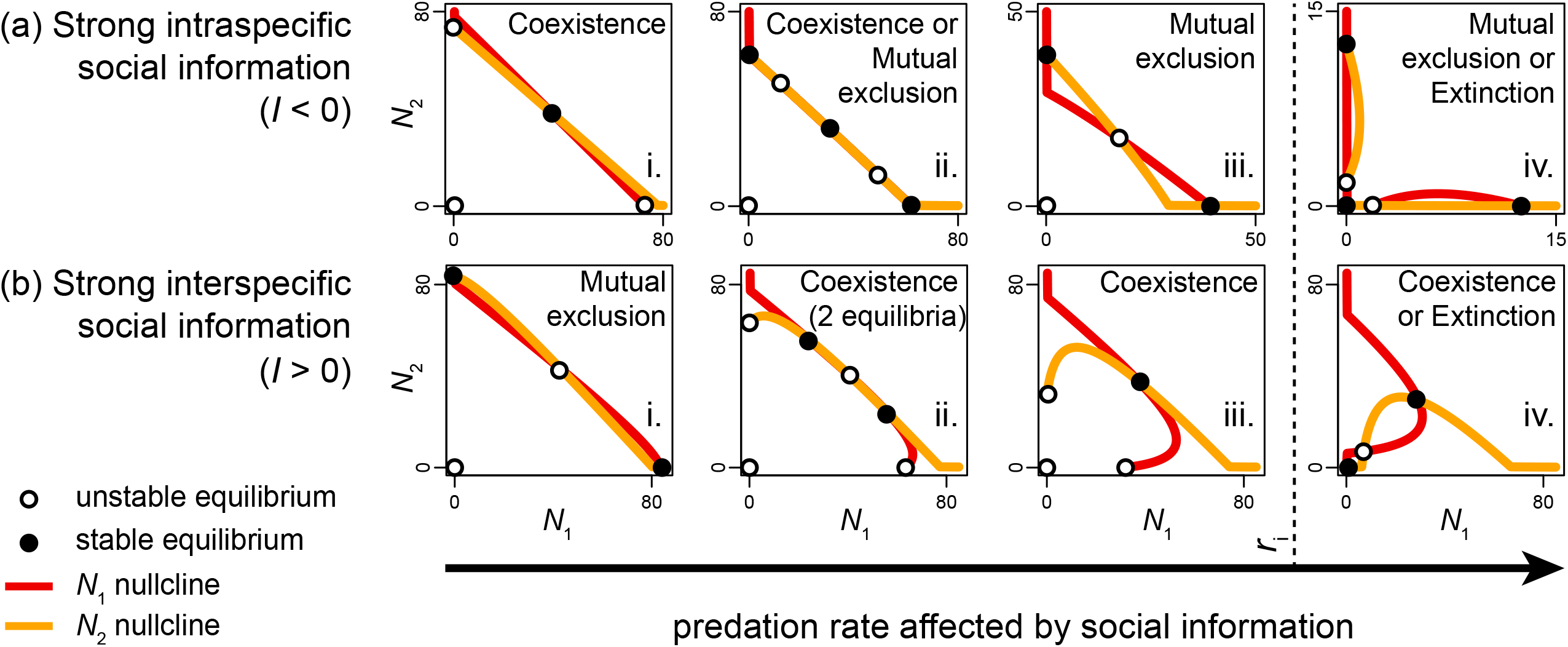
The interaction indices, which integrate the intraspecific and interspecific competition and social information, determine the coexistence-exclusion boundary when both forms of social information (intraspecific and interspecific) are weak (*I*_w_ = 0, A) and the persistence-extinction boundary when at least one form of social information is strong (*I*_s_ = 0, B). Parameter values: *r* = 1; *p* = 0.999999 (A), 1.002271 (B); *α* = 0.011 (A), 0.010 (B); 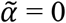 to 0.1; *b* = 0.010 (A), 0.011 (B); 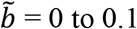 to 0.1.

When only one form of social information is strong, that form determines the sign of *I_s_*. If intraspecific social information is strong and interspecific social information is weak (i.e., 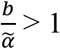 and 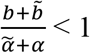), then *I^s^* is negative and social information promotes mutual exclusion as *p* increases (Fig. 4a). Alternatively, if interspecific social information is strong and intraspecific social information is weak (i.e., 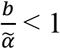 and 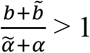), then *I_s_* is positive and social information promotes coexistence as *p_i_* increases (Fig. 4b, where Fig. 4biii is the specific case study in Gil et al 2018 Box 3 using a different functional form for the social information feedback).

**Fig. 4:**
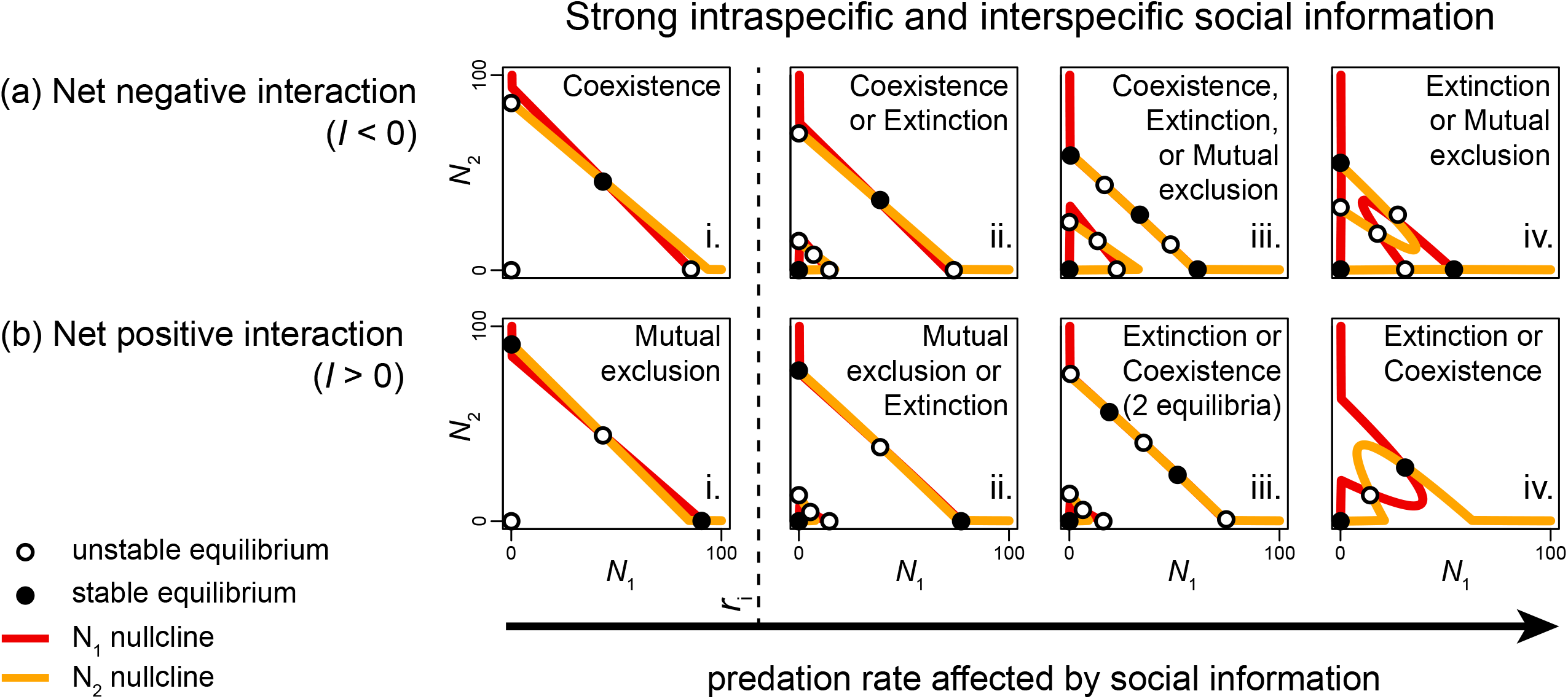
Phase plane plots of nullclines at which the population of each competing species exhibits zero net growth (*N*_1_ nullcline in red, *N*_2_ nullcline in yellow) when only one type of social information is strong in the system: intraspecific social information (a) or interspecific social information (b). Nullcline intersections mark equilibrium points (open points denote unstable equilibria; closed points denote stable equilibria). The type of social information that is strong determines the sign of interaction indices *I_s_* or *I_w_* (Eq. 4 & 5) and, thus, the progression of equilibrium outcomes that a system can experience, as the predation rate, *p*, which is affected by social information, increases (bottom *x*-axis). At the vertical dashed line, predation rate *p* exceeds the intrinsic growth rate *r* such that the extinction of all species would occur without social information. For *p* < *r*, coexistence would occur without social information in (a), and mutual exclusion would occur without social information in (b). Parameter values used are provided in the extended version of this figure (Appendix S3: Fig. S1).

With either form of strong social information, once the predation rate (*p*) exceeds the intrinsic rate of growth (*r*) of the two prey species, alternative stable states occur (as in the single-species model: Fig. 1), while both species would go extinct without social information. When *I_s_* is negative, there are three alternative stable states: each species persisting in isolation or mutual extinction, and coexistence can no longer occur (Fig. 4aiv). In other words, when predation is sufficiently high, strong intraspecific social information alone can cause competitors to become ‘obligate excluders’ (i.e., the only equilibria require competitive exclusion). Conversely, when *I_s_* is positive, single-species equilibria are eliminated, and there are two alternative stable states: species coexistence or extinction of all species (Fig. 4biv). Thus, strong interspecific social information alone can cause competitors to become obligate mutualists (i.e., the only equilibria require coexistence) at this critical level of predation.

Further increasing the predation rate results in the extinction of all species as the only outcome (see single-species analog in Appendix S2: Fig. S1c). Where exactly the extinction threshold for the two-species system occurs depends, again, on the sign of *I_s_*. When *I_s_* is negative, system-wide extinction occurs when the predation level *p* exceeds the critical predation level *p\** that we found for the single-species model (Eq. 3). When *I_s_* is positive, the critical predation level is

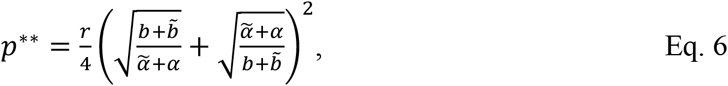

(see Appendix S3 for additional details).

When both forms of social information are strong (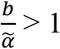 and 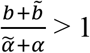), their opposing effects generate greater dynamical complexity with the introduction of additional alternative stable states. As before, when *I_s_* is negative (i.e., intraspecific social information is stronger than interspecific social information; Fig. 5a, Appendix S3: Fig. S2a) and as *p* increases, even a system with low interspecific competition will shift from outcomes at equilibrium that can include stable coexistence (Fig 5ai,ii & Appendix S3: Fig. S2ai-iii) to those that include only mutual exclusion or extinction (Fig. 5aiii,iv & Appendix S3: Fig. S2aiv-viii). Conversely, when *I_s_* is positive (i.e., intraspecific social information is weaker than interspecific social information; Fig. 5b, Appendix S3: Fig. S2b) and as *p* increases, even a system with high interspecific competition will shift from outcomes at equilibrium that can include mutual exclusion (Fig. 5bi,ii & Appendix S3: Fig. S2bi,ii) to those that include only coexistence or extinction (Fig. 5biii,iv & Appendix S3: Fig. S2biii.-viii). As before (Fig. 4), the system goes extinct when *I_s_* is negative and *p* exceeds *p*\* (Eq. 3; Appendix S3: Fig. S2aviii), or when *I_s_* is positive and *p* exceeds *p\*\** (Eq. 6; Appendix S3: Fig. S2bviii).

**Fig. 5:**
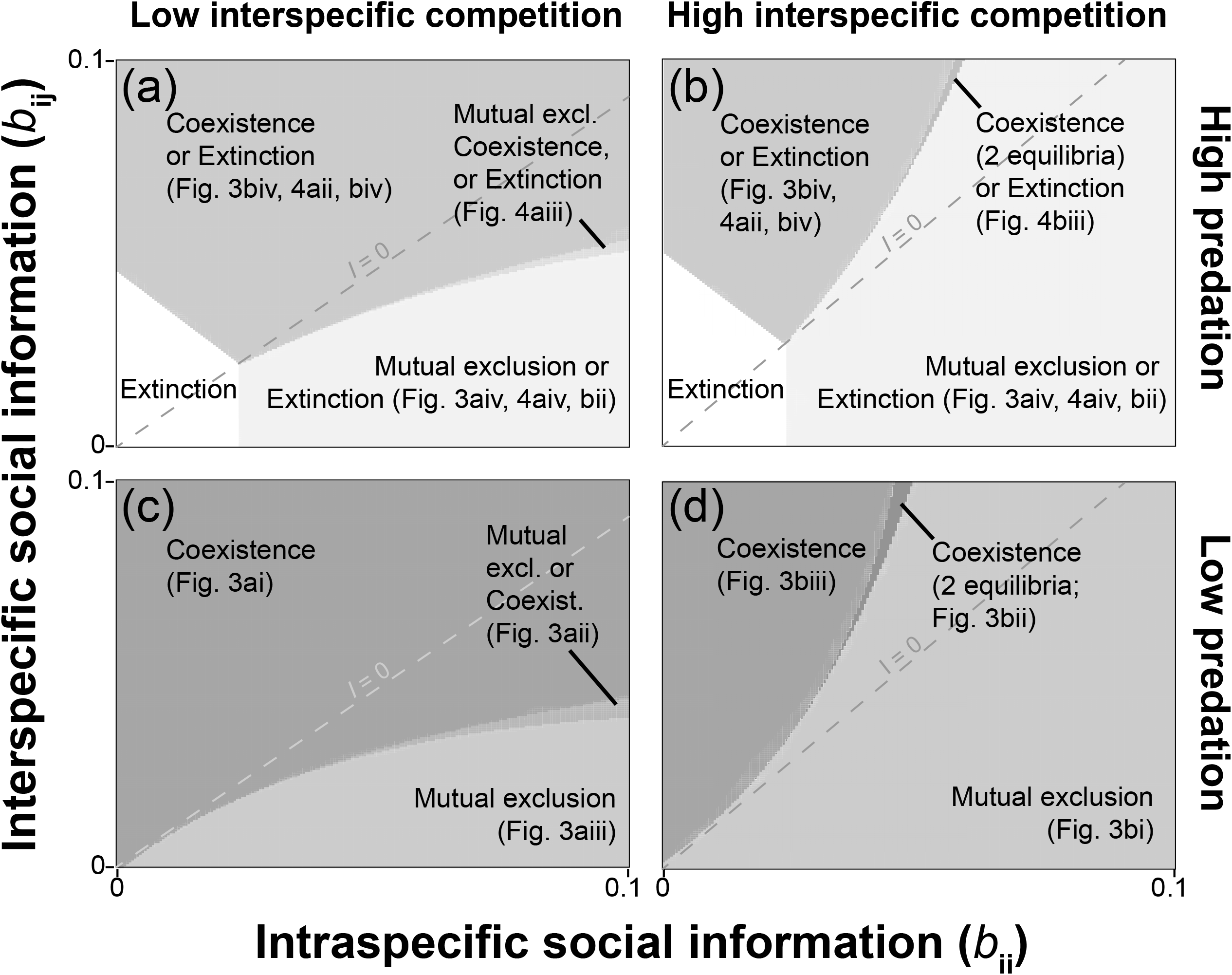
Phase plane plots of nullclines at which the population of each competing species exhibits zero net growth (*N*_1_ nullcline in red, *N*_2_ nullcline in yellow) when both intraspecific and interspecific social information are strong. Nullclines intersections mark equilibrium points (open points denote unstable equilibria; closed points denote stable equilibria). The sign of the net interaction index, *I_s_* or *I_w_* (calculated from the relative effects of social information and competition; Eq. 4 & 5), determines the progression of equilibrium outcomes that a system can experience, as the predation rate, *p*, which is affected by social information, increases (bottom x-axis). In (a), the interaction index is negative and promotes mutual exclusion. In (b), the interaction index is positive and promotes coexistence. At the vertical dashed line, *p* exceeds the intrinsic growth rate *r* such that the extinction of all species would occur without social information. For *p* < *r*, coexistence would occur without social information in (a), and mutual exclusion would occur without social information in (b). Parameter values used are provided in the extended version of this figure (Appendix S3: Fig. S2).

When predation rate, *p*, exceeds the intrinsic rate of growth, *r*, strong social information of both forms further increases the range of parameters where persistence and coexistence can occur (Fig. 5), in comparison to when only one form of social information is strong (Fig. 4). When *I_s_* is negative, coexistence equilibria do not vanish until predation rate exceeds *p\*\** (Eq. 6, the system-wide extinction threshold when only interspecific social information is strong), while single-species equilibria remain. When *I_s_* is positive, single-species equilibria do not vanish until predation rate exceeds *p\** (Eq. 3, the system-wide extinction threshold when only intraspecific social information is strong), while coexistence equilibria remain. Thus, when both forms of social information are strong, a greater diversity of prey community states are possible at high predation rates (Fig. 5), relative to cases when only one form of social information is strong (Fig. 4), or neither form is strong (in which case, system-wide extinction occurs when *p* > *r*, which is represented by the vertical dashed line in Fig. 4, 5; Appendix S3: Fig. S1, S2). Furthermore, while conditions that give rise to obligate excluders or obligate mutualists also emerge when both forms of social information are strong (Appendix S3: Fig. S2avi,vii, and Fig. S2bvi,vii), they generally do so over a narrower range of predation rates than when only one form of social information is strong.

### Context dependent effects of social information on competitive outcomes

Overall, qualitative shifts in competitive outcomes can occur under each of two conditions (Fig. 6, Appendix S3: Fig. S3): (1) predation exceeds population growth such that the positive effects of social information can rescue the system from extinction (Fig. 6, Appendix S3: S3a,b), or (2) strengths of intraspecific and interspecific social information are asymmetric in favor of an outcome that opposes that of competition. Regarding the second condition, high intraspecific social information can cause mutual exclusion under low interspecific competition [below the dashed line in Fig. 6c], or high interspecific social information can cause coexistence under high interspecific competition [above the dashed line in Fig. 6d]; Appendix S3: Fig. S3). As the strength of social information increases, greater asymmetries between intraspecific and interspecific social information are needed to qualitatively shift outcomes from expectations based on competition alone (see curved boundaries between coexistence and mutual exclusion: Fig. 6).

**Fig. 6.**
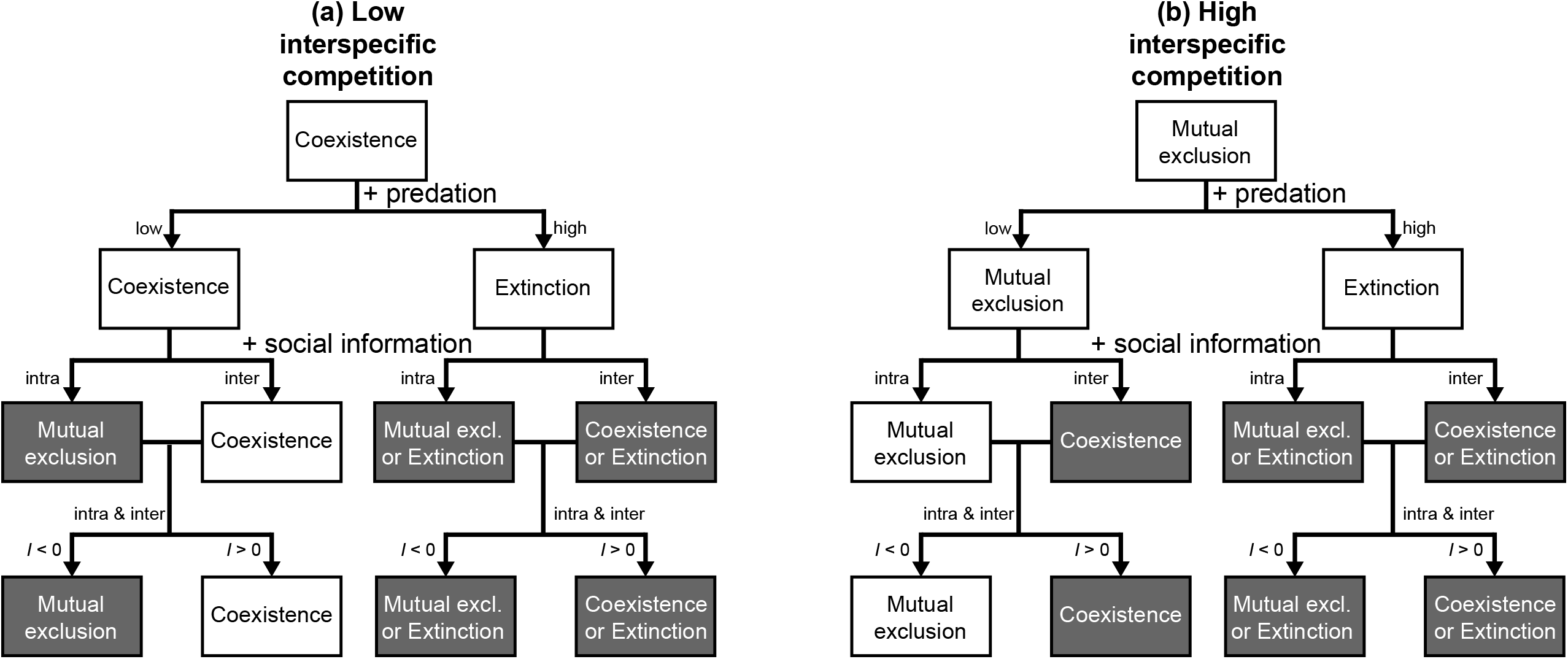
Social information drives qualitative shifts in the dynamics of competing species. The outcomes at equilibrium for competing populations respond to the relative strengths of social information types (intraspecific: x-axes; interspecific: y-axes). These responses depend on interspecific competition (columns: low (left; a, c): 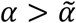, such that coexistence would occur without social information; and high (right; b, d): 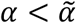, such that mutual exclusion would occur without social information), and predation (rows: high (top; a, b): *p* > *r*, such that system-wide extinction would occur without social information; and low (bottom; c, d): *p* < *r*). Furthermore, the direction in which social information influences the competitive outcome (towards coexistence or competitive exclusion), and therefore the suite of qualitatively distinct dynamics that the system can exhibit (e.g., Fig. 4a vs. 4b, Fig. 5a vs. 5b; Appendix S3: Fig. S1a vs. S1b, Fig. S2a vs. S2b), depends on the sign of the net interaction indices *I_s_, I_w_* (Eq. 4 & 5), where *I*_s_ = along the dashed line, *I*_s_ < 0 below the dashed line, and *I*_s_ > 0 above the dashed line. Parameter values: *r* = 1, *α* = 0.01, 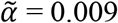 (left side; a, c) or 0.011 (right side; b, d), *p* = 1.2 (top row; a, b) or 0.9 (bottom row; c, d).

## Discussion

Our theoretical models reveal that the simple and ubiquitous use of social information by individual animals (e.g., using the alarm calls or flight responses of others to avoid danger) can scale up to qualitatively affect population and community outcomes. Specifically, our results indicate that by having positive effects on per capita population growth, even when net positive effects are restricted to low population densities, social information typically raises equilibrium population sizes and allows persistence, with Allee effects, when extinction would otherwise occur (due to either predation or interspecific competition in our models; Fig. 1, 6). These effects of social information on population and community stability arise because social information can decrease mortality and can give rise to critical population thresholds, and if a population or community falls below such a threshold it will have insufficient information from conspecifics or heterospecifics to grow and, thus, will be susceptible to sudden and rapid collapse (Fig. 1, 4, 5). Furthermore, we show that the community-level consequences of social information are strongly context dependent, where new metrics, the net interaction indices (Eq. 4 & 5), which measure relative strengths of intraspecific and interspecific social information and competition, determine the direction in which social information influences competition (towards coexistence or mutual exclusion) and, therefore, the suite of qualitative outcomes that are possible in a multi-species system (Fig. 3-6). Thus, social information can qualitatively change the long-term outcome of species interactions from mutual exclusion to coexistence or from coexistence to mutual exclusion, by allowing systems to overcome net effects of competition (i.e., intraspecific social information counters effects of intraspecific competition, and interspecific social information counters effects of interspecific competition; Fig. 6).

The types of qualitative differences in population and community dynamics with social information illustrated for two specific case studies in Gil et al (2018) can occur under a broad range of parameters, including a range of competitive interactions. As we develop this new theory, it is important to recognize the challenges of empirically measuring effects of social information in many natural systems. While notable work has been done to quantify behavioral effects of social information in the form of vocalizations in avian systems (Betts et al. 2008, Magrath et al. 2015), and this work has been used to inform demographic models of socially-enhanced resource acquisition (Schmidt et al. 2015, Schmidt 2017), social information is shared through more nuanced behaviors, such as movements, in many systems. Fortunately, recent advances in the collection of large, high-resolution datasets on individual behaviors in the wild, combined with probabilistic models, are able to reveal strong information-mediated behavioral effects that emerge from subtle individual movements (Strandburg-Peshkin et al. 2015, Gil and Hein 2017, Hein et al. 2018). These and other advances could aid in determining the functional form and parameter values of system-specific models that extrapolate these effects to their demographic consequences and test the theory we develop here.

### Single-species model

Our findings on the effects of social information on a single species expand upon the results of (Schmidt et al. 2015, Schmidt 2017), which showed that eavesdropping on breeding habitat quality can affect the dynamics and persistence of a population. Here, we model the use of intentional signals or unintentional cues about predators and show that social information could be a driver of positive density dependence and critical thresholds in relevant natural populations (Courchamp et al. 1999, Gascoigne and Lipcius 2004a, Suding and Hobbs 2009, Kelly et al. 2015).

Our findings further suggest that social information could serve as a stabilizing mechanism for predator-prey interactions: high levels of predation that would otherwise reduce prey populations to extinction and, consequently, threaten predator populations can be sustained when there is sufficient social information available to prey (Fig. 1, 3-6). Thus, social information about predators would be most important to the coexistence of predators and prey in systems in which predators can exert high pressure on individual prey populations (e.g., Van de Koppel et al. 2005, Sandin et al. 2008). Because we assume non-dynamic predators, demographic effects of social information in the face of such factors as dynamic predators and differential social information use across trophic levels remain important, unexplored topics for further research. Nonetheless, our models are representative of instances in which predator and prey populations are demographically decoupled (e.g., due to wide-dispersing or ranging predators; Hixon et al. 2002, Van de Koppel et al. 2005) and provide an important first step toward understanding how predation pressure can interact with effects of social information and competition to shape how populations of prey species grow, persist, and interact.

### Two-species model

Ecologists have long recognized that the qualitative nature of species interactions (negative, positive, neutral) are not static in space or time, but vary on a continuum in nature (Bronstein 1994, 2001). Understanding the context dependence of species interactions that shape fundamental rates at the population, community and ecosystem levels remains an open but pressing challenge in the discipline of ecology (Agrawal et al. 2007). Here, we provide theory that shows that a common driver of animal behavior, social information, can be a powerful force that shapes the strength or sign of species interactions.

We show that the fate of competing species can be determined not by the relative strengths of intraspecific and interspecific competition, per se (as we conventionally expect), but instead by the interplay between competition and social information (Fig. 4, 5), Further, we show that because positive effects of social information can strongly affect demographics at low to moderate population densities, while negative effects of competition are strongest at high densities, these opposing effects do not simply cancel one another out but instead interact to give rise to a range of stable population states. For example, we show that strong intraspecific relative to interspecific competition can fail to cause long-term coexistence, as we would otherwise expect, if the effect of intraspecific social information is stronger than the effect of interspecific social information (Fig. 4a, 5a). In nature, this scenario could result when predators of competing species differ to enough of a degree that social information about predators is more valuable when it comes from conspecifics than when it comes from heterospecifics, or in the extreme case that intraspecific social information, alone, is of value (Seppänen et al. 2007). Conversely, we show that weak intraspecific competition relative to interspecific competition can fail to drive a system to long-term mutual exclusion, as we would otherwise expect, if the effect of intraspecific social information is weaker than the effect of interspecific social information (Fig. 4b, 5b). In nature, such a scenario could result when individuals are distributed in space such that they are more likely to be proximate to (and, thus, privy to information from) heterospecifics than conspecifics (e.g., in mixed-species bird flocks; Graves and Gotelli 1993, Greenberg 2000, Templeton and Greene 2007, Martínez et al. 2018). Effects of interspecific (relative to intraspecific) social information can be further enhanced by phenotypic differences among species, allowing some species heightened sensory abilities and/or more effective means of transmitting information (Seppänen et al. 2007, Goodale et al. 2010). In either case, surrounding species can come to rely upon such information producers; for example, various bird species are highly responsive to even the nuances of alarm calls from keystone informant species (Templeton et al. 2005, Templeton and Greene 2007), and similarly, zebras respond strongly to simple movements of giraffes (i.e., body postures directed at predators), which possess a much higher vantage point to spot shared predators (Schmitt et al. 2016). Thus, while we may typically expect the strength of the effect of social information on prey demographics to be more pronounced in individuals that more frequently aggregate (e.g., in cohesive flocks, herds or schools), highly useful information (e.g., that prevents predation) could have strong demographic effects even when it is received infrequently.

To simplify our presentation and for analytical tractability (Appendix S3), we primarily focus on symmetrically competing populations. However, the same principles revealed above apply to cases when competing populations exhibit differences in competitive ability: social information counters effects of competition, within or between species, and, consequently, can tip the scales in favor of competitively inferior species or strengthen the dominance of competitively superior species (Appendix S4: Fig. S1). Furthermore, we assume that competing species share a generalist predator and that social information reduces encounter rates with this predator. However, a specialist predator would create dissimilarities in the value of social information between species, such that the prey species that is preferred by the predator would exhibit a stronger positive response to social information than the less-preferred prey species, and this can affect the long-term outcome for both prey populations (Appendix S4: Fig. S2). It is also true that when social information enhances resource acquisition (Dall et al. 2005, Goodale et al. 2010), instead of or in addition to enhancing predator avoidance, it could exacerbate resource or interference competition in certain contexts (Gil et al. 2017). Consequently, other factors, such as the abundance and distribution of resources, could strongly influence the overall effect of social information on competitive outcomes. In summary, our models show that when net effects of social information exceed and oppose net effects of competition, social information can affect the qualitative outcome at equilibrium, but there remains vast opportunity to expand our framework to incorporate specific features of natural history of various systems.

### Conclusions

Mounting empirical and theoretical evidence suggests that social information, a ubiquitous driver of animal behavior and fitness (Goodale et al. 2010, Magrath et al. 2015), can play a significant role in the ecology of natural systems (Gil et al. 2018). Yet, our paper is the first to our knowledge to formalize the inclusion of social information in both single and multi-species population models and to rigorously characterize the demographic effects thereof. Our study provides an important step in our understanding of the potential for social information to underlie the persistence, coexistence and diversity of species across systems.

Our study also highlights a multifaceted potential impact of social information on expectations for conservation management. Social information can raise population size and allow persistence that we would not otherwise expect (e.g., due to high predation and/or competition); however, under conditions of high predation, social information can cause putatively common critical population thresholds (Courchamp et al. 1999, Gascoigne and Lipcius 2004a, Suding and Hobbs 2009, Kelly et al. 2015) that, if not identified by resource managers, can lead to unrealized risks of sudden population collapse (Holt 2007). The probability of such information-mediated local extinctions could increase with demographic or environmental stochasticity (Gilpin and Soulé 1986, Lande 1998). Furthermore, the demographic effects of social information that we reveal suggest that environmental changes that simply inhibit social cueing/signaling among individuals (e.g., anthropogenic increases in turbidity or disruption of chemical cues in aquatic systems (Kimbell and Morrell 2015, Chivers et al. 2016), urban noise masking auditory signals in terrestrial systems (Patricelli and Blickley 2006)) could drive unexpected changes to extinction risk, the outcome of competition, and ultimately the community state (Holt 2007). Furthermore, by changing the expected community structure as it depends on competitive and predation rates, social information could affect the expected ecological outcome of an invasive predator or competitor. The directionality of this effect on expected invasiveness and invasive impact will inevitably depend on the interplay between competition and social information, as described above. Consequently, our findings point to social information as an important factor that could affect how we conserve and manage natural resources, particularly for endangered species at small population sizes where social information is more likely to influence demographic rates.

## Acknowledgements

We thank Andrew Hein, Andy Sih and Orr Spiegel for helpful feedback in early discussions of this model. This research was funded by a National Science Foundation Postdoctoral Research Fellowship awarded to MAG.

## Supporting Information. Appendix S1: Mechanistic derivation and re-parameterizations

Michael A. Gil, Marrisa L. Baskett, and Sebastian J. Schreiber. 2019. Social information drives ecological outcomes among competing species. Submitted to *Ecology*

In this Appendix, we provide a mechanistic derivation of the predation term for the models in the main text, and describe how these models can account for a minimal level of predation as well as a type II functional response. We do the re-parameterizations separately to minimize the amount of notation.

### A mechanistic derivation

Here, we show how the functional form of our predation term in the two species model can be derived from first principles. For illustrative purposes, assume that the population of species *i* consists of individuals that are vulnerable to predation (with density *V_i_*) and informed individuals (with density *I_i_*) that are invulnerable to predation; in the next section, we describe how to account for a minimal level of predation on all individuals (i.e., even informed individuals are subject to some level of predation). We have that *N_i_* = *V_i_* + *I_i_*. Invulnerable individuals return to being vulnerable at a characteristic rate *γ_i_* corresponding to their tendency to return to less informed behavior. Alternatively, vulnerable individuals by interacting with intra- and inter-specific individuals gain social information about the dangers of predation. If the movement between these behavioral stages occurs at a faster time scale than changes in population densities, then we can describe shifts between these behavioral states as

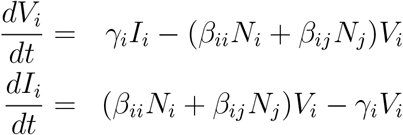

where *β_ii_, β_ij_* are the per-capita influence of intra- and interspecific individuals (via social information) on switching from vulnerable to informed. Over the faster behavioral time scale, these behavioral dynamics converge to a unique equilibrium which satisfies

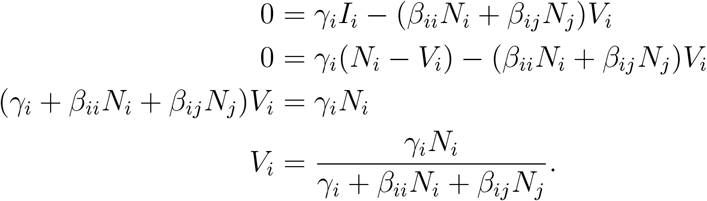

Dividing the numerator and denominator of the final expression by *γ_i_*, we get

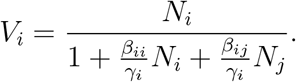

As only vulnerable individuals experience predation, the net predation rate on species *i* equals

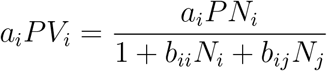

where *b_ii_* = *β_ii_/γ_i_, b_ij_* = *β_ij_/γ_i_*, and *P* is the predator density. This functional form is used in all of the models in the main text.

### Accounting for minimal predation

First, we show how a model which includes a minimal level of predation 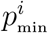, for each species *i* = 1, 2, that occurs irregardless of social information corresponds to a re-parameterization of the model presented in the Models and Methods. As in the model presented in the main manuscript (single-species model: Eq. 1, two-species model: Eq. 2), *r_i_* is the intrinsic growth rate of species *i*, *α_ij_* is the per-capita competition coefficient for the effect of species *j* on species *i, p_i_* is the additional maximal predation level that occurs when there is no social information, and *b_ij_* determines the per-capita reduction of predation on species *i* due to social information from species *j*. This model is given by

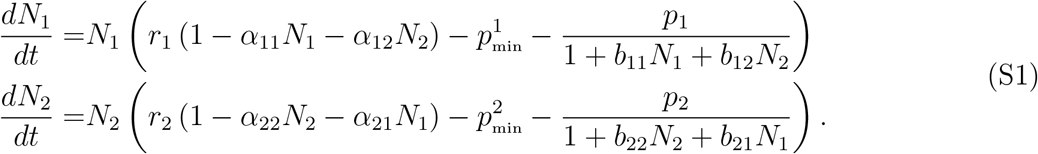

Assume 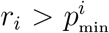 for *i* = 1, 2. Define 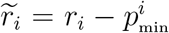 and 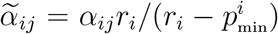. Then 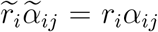 and we get

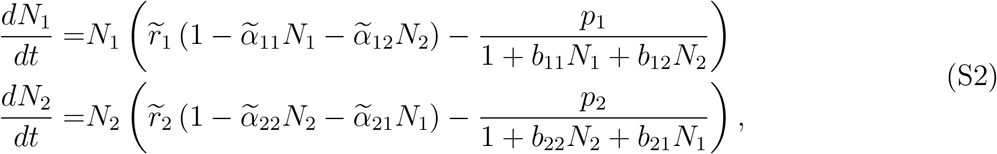

which has the same form of the model shown in Eq. 2 in the main text.

### Accounting for a type II functional response

To account for a type II functional response for the generalist predator, we begin with the single species model and then discuss the two species model. For the single species model, we shall show that it is equivalent, via re-parameterization, to the model studied in the main manuscript. For the two species model, we shall show that it is equivalent to the model presented in the main text under special circumstances that allow some of the analysis to extend to the two species model with a type II functional response.

For the single species model, let *h* be the handling time of the predator. The attack rate of the predator is a decreasing function of intraspecific social information *a*/(1 + *bN*) where *a* is the maximal attack rate of the predator. Under these assumptions the functional response of the predator is given by

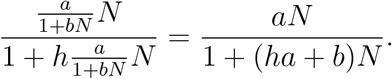

If *P* is the total density of the generalist predator, then the model becomes

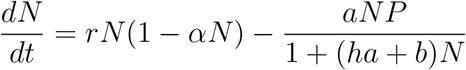

Setting 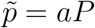 and 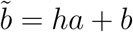, we get

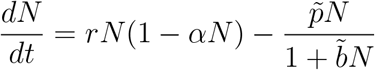

that is equivalent to Eq. 1 in the main text.

Now, consider the two species model where the generalist predator has a handling time *h_i_* on species *i*, and an attack rate *a_i_*/(1 + *b_ii_N_i_* + *b_ij_N_j_*) on species *i*. Then its functional response with respect to species 1, for example, is

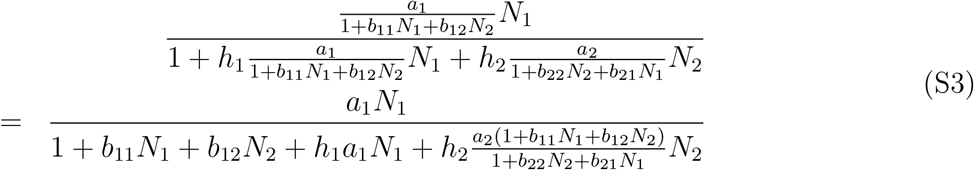

Unlike the functional response with a single prey species, the term 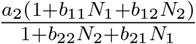 in the denominator of equation (S3) implies that this expression for the predator’s functional response does not always simplify to an expression equivalent to the functional response in Eq. 2 in the main text. Two special cases where it does are as follows. First, if inter- and intraspecific social information terms are equal (i.e. *b*_11_ = *b*_22_ = *b*_12_ = *b*_21_), then the ratio 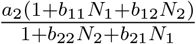 reduces to *a*_2_ and this functional response is equivalent to the one presented in the main text. Second, if there is symmetry in the social information cues (i.e. *α*_11_ = *α*_22_ and *α*_12_ = *α*_21_) and both species have equal densities (i.e. *N*_1_ = *N*_2_ =: *N*), then the ratio 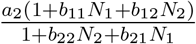 also reduces to *a*_2_. It follows that the bifurcation analysis of the symmetric model along the *N*_1_, *N*_2_, *N*_1_ = *N*_2_ axes in Supplementary Information S3 still applies where *h*_1_ = *h*_2_, *a*_1_ = *a*_2_, *p_i_* is replaced with *a_i_P, b_ii_* is replaced with *h_i_a_i_* + *b_ii_*, and *b_ij_* is replaced with *h_i_a_i_* + *b_ij_*. In particular, the interaction index in this case is given by 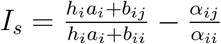.

## Supporting Information. Appendix S2: Analsysis of single species model

In this Appendix, we analyze the single-species model presented in the main text and numerically explore an alternative formulation of the model. Although the main model is mathematically equivalent to the model of Noy-Meir [1975], our analysis of the two possible types of bifurcations and the first order approximation of the positive equilibrium are novel.

### The bifurcation analysis

Recall, the single species model is

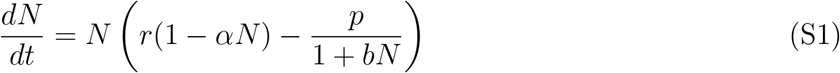

where *N* is the population density, *r* is the intrinsic rate of growth, *α* is the strength of intraspecific competition, *p* corresponds to the maximal predation level which can be reduced to near zero by social information, and *b* is the per-capita reduction in predation due to intraspecific social information. Positive equilibria of (S1) must satisfy

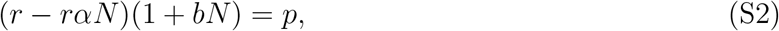

or equivalently

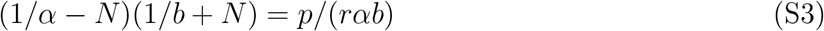

As the left hand side is a quadratic with roots at 1/*α* and −1/*b* and a maximum of (1/*α* + 1/*b*)^2^/4 at (1/*α* − 1/*b*)/2, we get two possible sequences of bifurcations as *p* increases from zero to infinity (Figure S1):

*Weak social information (b/α < 1)*: If *p* < *r*, then the population persists at a globally stable positive equilibrium *n*\*. If *p* > *r*, then the population goes extinct as *n* = 0 is a globally stable equilibrium.

*Strong social information:* Assume *b/α* > 1. If *p* < *r*, then the population persists at a globally stable feasible equilibrium *n*\*. If 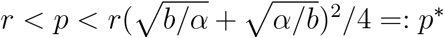(the same as Eq.3 in the main text), there are two feasible equilibria *n*_*_ < *n*\*, such that initial conditions below *n*\* go to extinction and initial conditions above *n*_*_ converge to the stable equilibrium *n*\*. If *p* > *p*\*, then the population goes asymptotically extinct for all initial conditions as *n* = 0 is globally stable.

Note that the critical value *p*\* is proportional to *r*, and increases from the value of *r* up to ∞ as *b/α* goes from 1 to ∞. Namely, the stronger the social information, the higher level of predation that the population can withstand.

### Effects of social information on equilibrium densities

To understand the effect of social information *b* on the non-zero equilibria of the model, let *R* denote the per-capita growth rate of the population i.e.

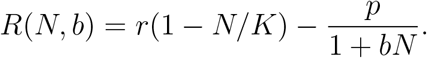

**Figure S1:**
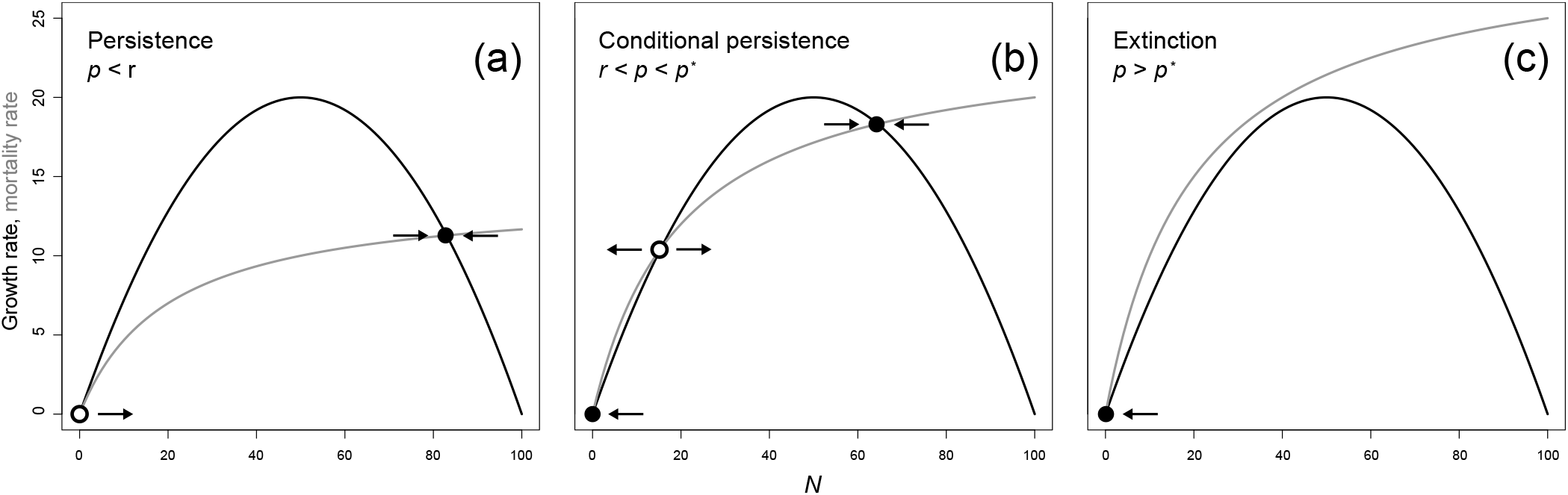
Social information can have quantitative or qualitative effects on a population. If social information is absent (i.e., *b* = 0 in Eq. 1), the prey mortality rate increases linearly with the prey population, with the slope = *p*. However, when social information is present, it causes the prey mortality rate (grey curves, with initial slope = *p*) to decelerate with prey density (*N*, *x*-axis), allowing for either a greater population size at equilibrium (a; a quantitative effect) or conditional persistence when we would otherwise expect extinction (b; a qualitative effect), if *p* < *p*\* (Eq. 3 in the main text). However, when *p* > *p*\*, social information has no effect (i.e., the population goes extinct whether or not social information is available; c).

Then the growth rate is *G*(*N, b*) = *NR*(*N, b*). A positive equilibrium density *N*\*(*b*) for this model satisfies *R*(*N*\*(*b*), *b*) = 0 and is stable if

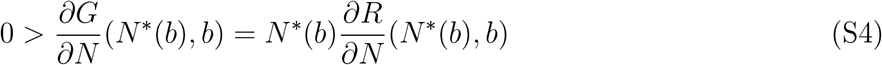

and unstable if

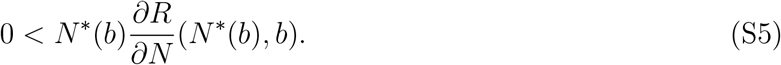

To understand how *N*\*(*b*) varies with *b*, we can implicitly differentiate with respect to *b*

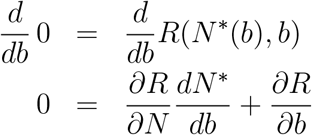

Therefore, whenever 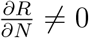,

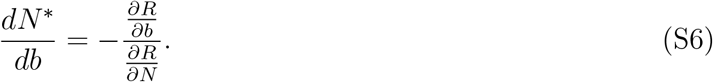

As

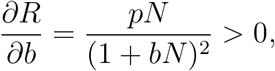

equations (S4)–(S6) imply that 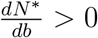 when *N*\* is stable and 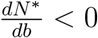 when *N*\* is unstable.

### An alternative functional form of the model

As an alternative functional form of the effect of social information on predation, we also consider an inverse normal function to model cases in which reductions in per capita mortality due to social information manifest at low densities but are completely negated at higher densities (e.g., due to false alarms and/or occlusion of information [Rosenthal et al., 2015]). In this form, *p_max_* sets the effect of social information on reducing mortality due to predation (analogous to *p* in Eq. 1 in main text), b controls the strength of the effect of social information and the (symmetric) strength of compensation, and N* is the population size at which mortality due to predation is minimized by social information, such that the effective predation rate is determined by the second term in the model:

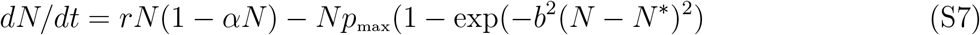

**Figure S2:**
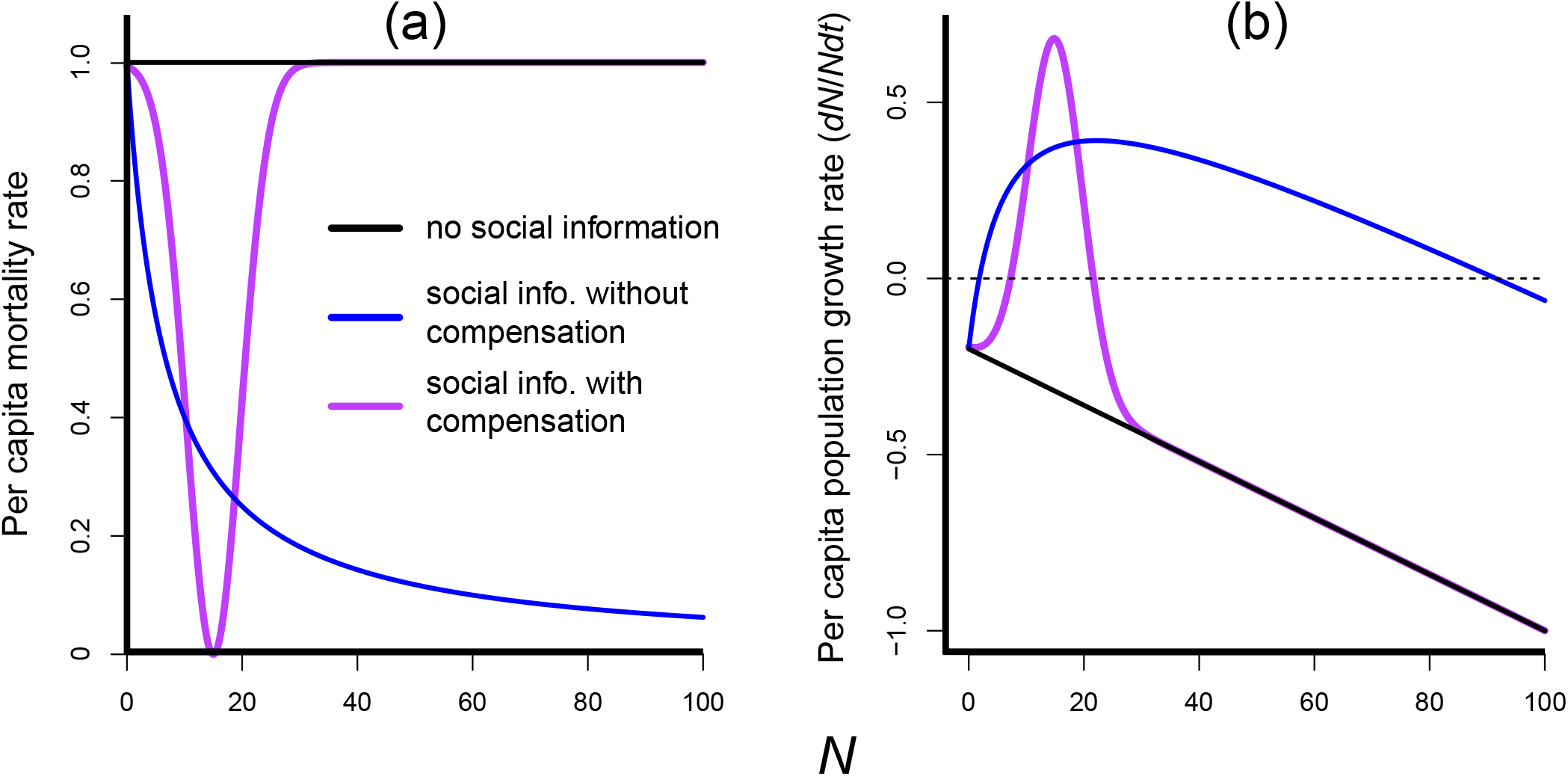
Effects of social information on population dynamics are robust across functional forms. Inclusion of effects of social information on per capita mortality due to predation in two distinct functional forms (a) gives rise to positive density dependence in the per capita population growth rate (b) and expands, relative to the logistic model with predation (black line), the conditions under which a population can persist. In particular, when predation rate exceeds population growth rate (*p* > *r* for the model with social information [blue curves] or *p*_max_ > *r* for the model with social information and compensation [purple curves]), as it does in (b), social information can give rise to alternative stable states. Thus, social information can prevent population collapse if population size does not fall below an unstable equilibrium that represents a critical threshold (where the blue and purple curves first intersect the dotted line at *y* = 0 in (b); a ‘strong Allee effect’. For these calculations, we set *r* = 0.8, *α* = 0.01, *p* = *p*_max_ = 1, *N*\* = 15, and *b* = 0 (Eq. 1 and Eq. 2–8 in the main text) for ‘no social information’, or *b* = 0.15 for both models with social information. Note that the inverse normal functional form (purple curve) has the additional property of bistability between low and high population sizes over a narrow range of *p*, when *p* > *r*.

Note that complete compensation for effects of social information is likely less common than cases of partial compensation (i.e., when benefits of social information are only partially negated at higher densities) [Kenward, 1978, Seppänen et al., 2007, Jackson et al., 2008, Lister, 2014, Berdahl et al., 2016]. Thus, the functional form of predation in Eq. S7 can be considered a lower bound of the demographic consequences of social information. Nonetheless, this functional form with compensation drives the same qualitative pattern as the monotonic form (without compensation; Eq. 1; Fig. S2): it allows for a greater carrying capacity of the population (relative to the logistic + predation model) and can prevent extinction, when predation exceeds the intrinsic rate of growth (*p*_max_ > *r*; Fig. S2).

## Supporting Information. Appendix S3: Analysis of two-species model

In this Appendix, we analyze the two-species competition model presented in the main text. Recall, this model is given by

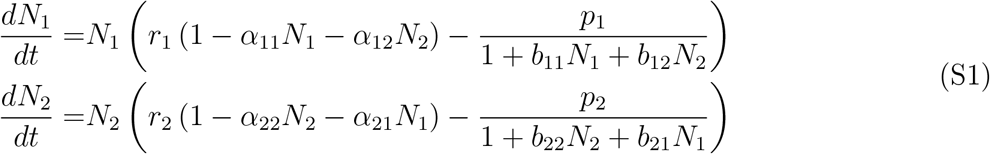

where *r_i_* is the intrinsic rate of growth of species *i*, *α_ij_* is the strength of the competitive effect of species *j* on species *i*, *p*_*i*_ is the maximal predation level in the absence of social information, and *b_ij_* is the per-capita effect of social information from species *j* to species *i*.

### Bifurcation Analysis of the Symmetric Case

Consider the symmetric case i.e. 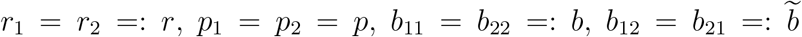, *α*_11_ = *α*_22_ =: *α*, and 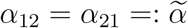.

Under these assumptions, there is an invariant line *N*_1_ = *N*_2_ =: *N* for the dynamics, and the dynamics on the line are given by

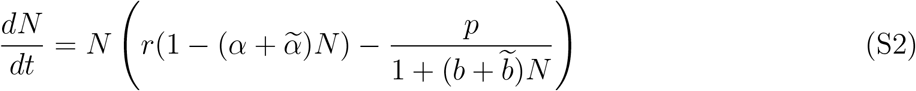

Our analysis of the symmetric model is divided into two parts. First, we identify when increasing *p* can cause the non-zero nullclines to cross on the single species axis. When intraspecific information is weak (i.e. *b/α* < 1 cf. the analysis in Appendix S2), this bifurcation corresponds to the system switching from bistability (i.e. both single species equilibria are stable) to coexistence in the sense of mutual invasibility (i.e. both single species equilibria are unstable), or vice-versa. Second, we study the equilibrium structure on the single species axes and the *N*_1_ = *N*_2_ axis. Together, these analyses provide the analytical scaffolding for the results presented in the main text. These analyses, however, do not address the structure of the asymmetric equilibria, i.e. pairs of equilibria of the form (*N*_1_; *N*_2_) = (*a, b*) and (*N*_1_; *N*_2_) = (*b, a*) with *a* > *b* > 0.

### Bifurcations along the single species and symmetric axes

As equation (S2) is the same as the single species model but with *α* replaced by 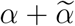 and *b* replaced by 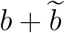, we can classify the bifurcations as *p* increases into 4 types. For this classification, we define two critical predation levels:

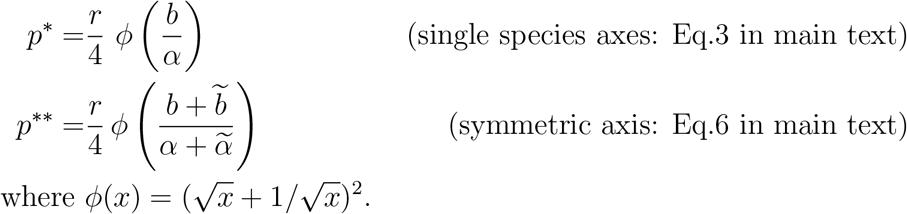

As *ϕ*(*x*) in an increasing function for *x* ≥ 1, *p*\*\* > *p*\* when 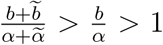 and, conversely, *p*\*\* < *p*\* when 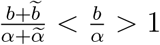. Furthermore, notice that

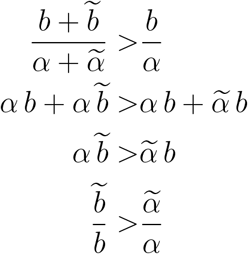

Thus, the interaction index 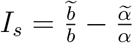 defined in Eqn. 5 in the main text is positive if and only if 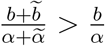, and *I_s_* < 0 if and only if 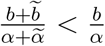. Based on these observation, we get the following cases:

*Weak intra and interspecific information* (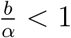 *and* 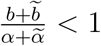): If *r* > *p*, then there are positive equilibria on each of these axes. If *r* < *p*, then there are no positive equilibria on these axes and extinction occurs for all initial conditions. As in this case, there is at most one positive equilibrium on each of the single species axes, our earlier analysis implies that the sign of interaction index *I_w_* determines whether predation can shift the system from coexistence to bistability (*I_w_* < 0) or vice-versa (*I_w_* > 0).
*Strong intra and weak interspecific information (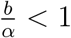 and 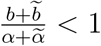):* As one increases *p*, one goes from a unique positive equilibrium on each axis (when *p* < *r*), to having no equilibria on the symmetric species axes and two positive equilibria on each of the single species axes (when *r* < *p* < *p*\*), to finally having no positive equilibria on any axis (when *p* > *p*\*). Notice that *I_s_* < 0 in this case as 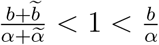. See Figure S1a.
*Weak intra and strong interspecific information (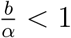 and 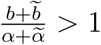):* As one increases *p*, one goes from a unique positive equilibrium on each axis (when *p* < *r*), to having no equilibria on the single species axes and two equilibria on the symmetric axis (when *r* < *p* < *p*\*\*), to finally having no positive equilibria on any axis (when *p* > *p*\*\*). Notice that *I_s_* > 0 in this case as 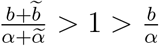. See Figure S1b.
*Strong intra and interspecific information (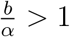 and 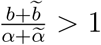):* If *p* < *r*, then there is a unique positive equilibrium on each axis. If *r* < *p* < min{*p*\*, *p*\*\*}, there are two positive equilibria on all three axes. If *p* > *p*\*, then all initial conditions on the single species axis go to extinction. If *p* > *p*\*\*, then all initial conditions on symmetric two species axis go to extinction. Depending on whether *p*\* > *p*\*\* or *p*\* < *p*\*\* one gets different orders of the bifurcations. If *I_s_* < 0, then *p*\* > *p*\*\* and one first loses the positive equilibria on the symmetric axis followed by the positive equilibria on the single species axes (Fig. S2a). If *I_s_* > 0, then *p*\*\* > *p*\* and one first loses the positive equilibria on the singles species axes and then the positive equilibria on the symmetric axis (Fig. S2b).

### Bifurcations from coexistence to bistability and vice versa

The invasion growth rates change sign as one increases *p* if and only if there is a *p* value at which the *N*_1_ and *N*_2_ nullclines intersect at the same point on the *N*_1_ axis (by symmetry, this intersection also occurs on the *N*_2_ axis). When such an intersection occurs, one has that *N*_1_ = *x* satisfies

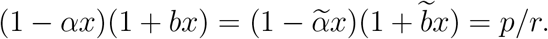

Equivalently,

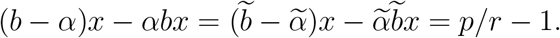

From the first equality, we get

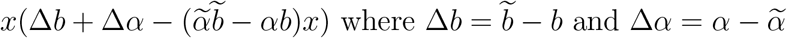

and either *x* = 0 (in which case *p* = *r*) or

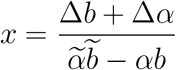

A nullcline crossing at *N*_1_ = *x* is only of interest if *x* > 0. Hence, we get two cases. First, if 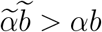, then *x* is positive if and only if Δ*b* + Δ*α* > 0. Notice that the quantity Δ*b* + Δ*α* corresponds to the interaction index 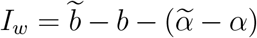 presented in Eq. 4 of the main text. Second, if 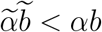, then x is positive if and only if *I_w_* < 0.

These observations have two implications. Recall that Δ*b* > 0 means interspecific information is greater than intraspecific information, and Δ*α* > 0 means that intraspecific competition is greater than interspecific competition. If Δ*α* < 0 (i.e. bistability in the absence of predation) and Δ*b* > 0, then predation can reverse the sign of the invasion growth rates (i.e. make them positive and thus allow for coexistence) only if *I_w_* > 0. Second, if Δ*a* > 0 (i.e. coexistence in the absence of predation) and Δ*b* < 0, then the sign of the invasion growth rates are reversed (i.e. both negative resulting in bistability) only if *I_w_* < 0.

**Figure S1:**
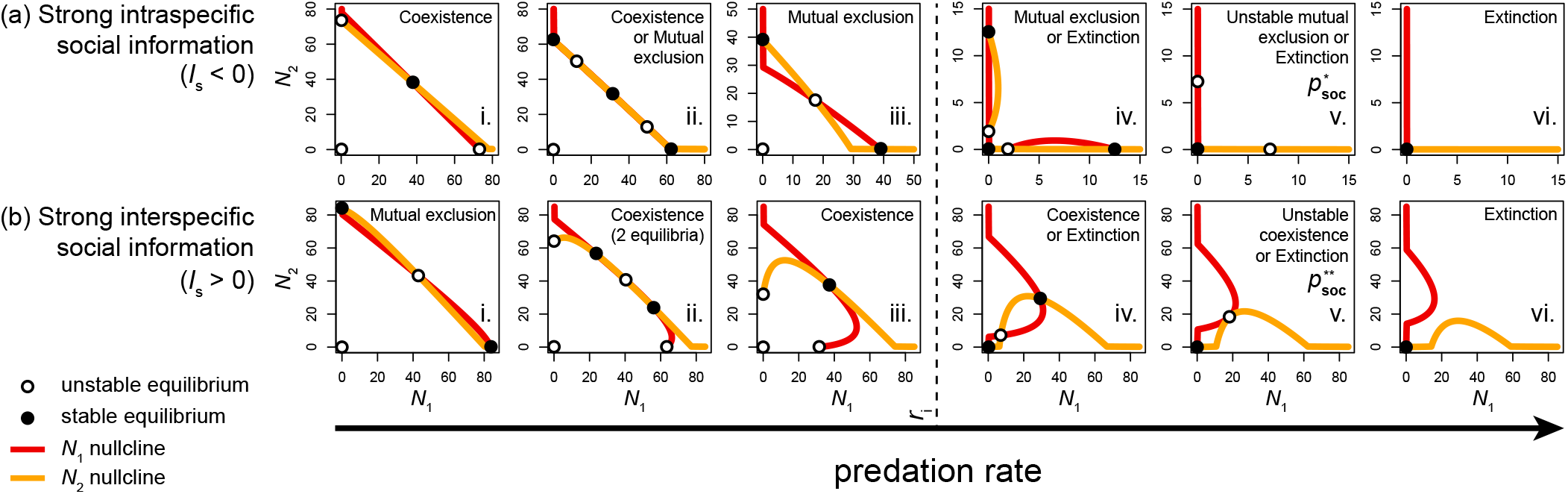
Phase plane plots of nullclines at which the population of each competing species exhibits zero net growth (*N*_1_ nullcline in red, *N*_2_ nullcline in yellow). Where nullclines intersect mark equilibrium points (open points denote unstable equilibria; closed points denote stable equilibria). Qualitatively distinct dynamics emerge when only one type of social information is strong in the system, intraspecific social information (a) or interspecific social information (b). The type of social information that is strong determines the sign of *I_s_* (Eq. 5 in main text) and, thus, the progression of equilibrium outcome that a system can experience, as the predation rate affected by social information, *p*, increases (bottom x-axis). When only intraspecific social information is strong (a), *p*\* (Eq. 3 in main text) marks the threshold predation level (av), above which the populations go extinct (avi), and when only interspecific social information is strong (b), *p*\*\* (Eq. 5 in main text) marks the threshold predation level (bv) above which the populations go extinct (bvi). Parameter values: *r* = 1; (a): *α* = 0.013; 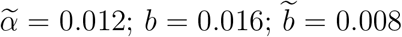; (b): *α* = 0.01; 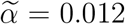; *b* = 0.01; 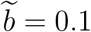; *p* = 0.1 (ai), 0.38 (aii), 0.8 (aiii), 1.004 (aiv), 1.01081T (av), 1.1 (avi), 0.3 (bi), 0.6 (bii), 0.9 (biii), 1.5 (biv), 1.8 (bv), 2 (bvi). Based on these parameters and Eq. 5, *I_s_* = −0.42 (a) or 8.8 (b).

**Figure S2:**
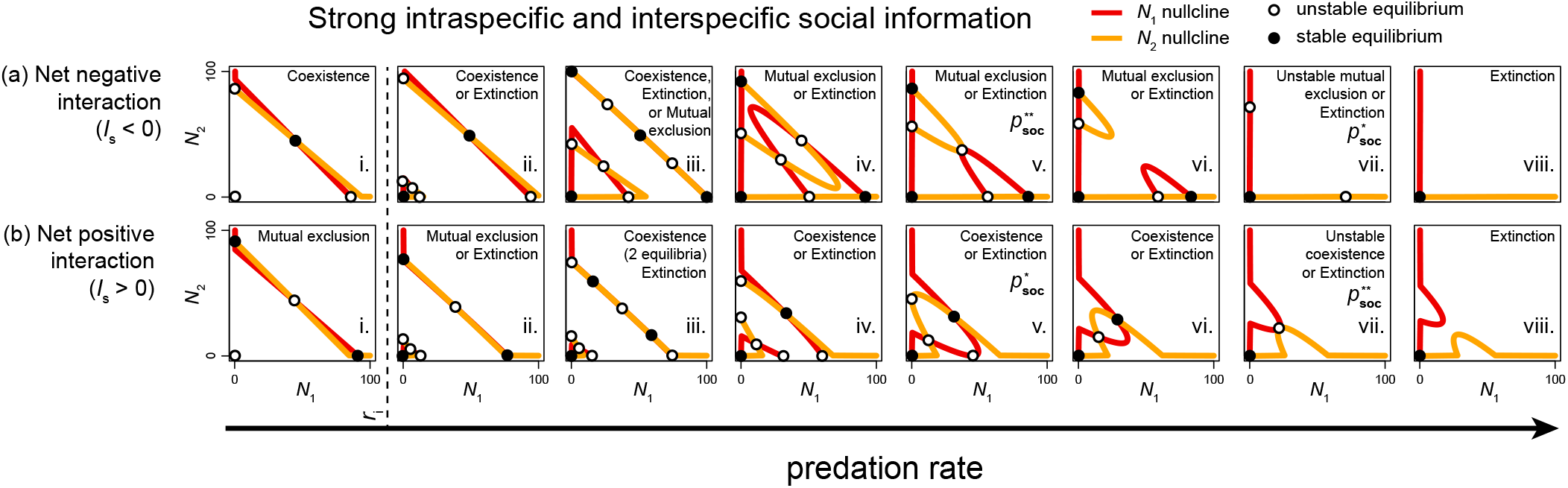
Phase plane plots of nullclines at which the population of each competing species exhibits zero net growth (*N*_1_ nullcline in red, *N*_2_ nullcline in yellow). Where nullclines intersect mark equilibrium points (open points denote unstable equilibria; closed points denote stable equilibria). Qualitatively distinct dynamics emerge when both intraspecific and interspecific social information are strong in the system. Under these conditions, as before (Fig. S1), it is whether the sign of the net interaction index, *I_s_* (calculated from the relative effects of social information and competition; Eq. 5 of main text), is negative (a) or positive (b) that determines the progression of equilibrium outcomes that a system can experience, as the predation rate affected by social information, p, increases (bottom x-axis). Here, equilibria on the single species axes vanish once predation exceeds *p*\* (avii, bv; Eq. 3 of main text), and equilibria on the symmetric two-species axis (the 1:1 line) vanish once predation exceeds *p*\*\* (av, bvii; Eq. 6 of main text). Parameter values: *r* = 1; (a): *α* = 0.011; 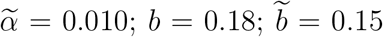; (b): *α* = 0.010; 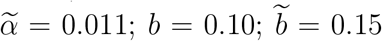; *p* = 0.9 (ai), 2 (aii), 2.15 (aiii), 2.T (aiv), 3.025 (av), 3.2 (avi), 3.49T19 (avii), 3.6 (aviii), 0.9 (bi), 3 (bii), 4 (biii), 4.35 (biv), 4.444481 (bv), 4.5 (bvi), 4.60618T (bvii), 4.7 (bviii). Based on these parameters and Eq. 5 of main text, *I_s_* = −0.08 (a) or 1.39 (b).

**Figure S3:**
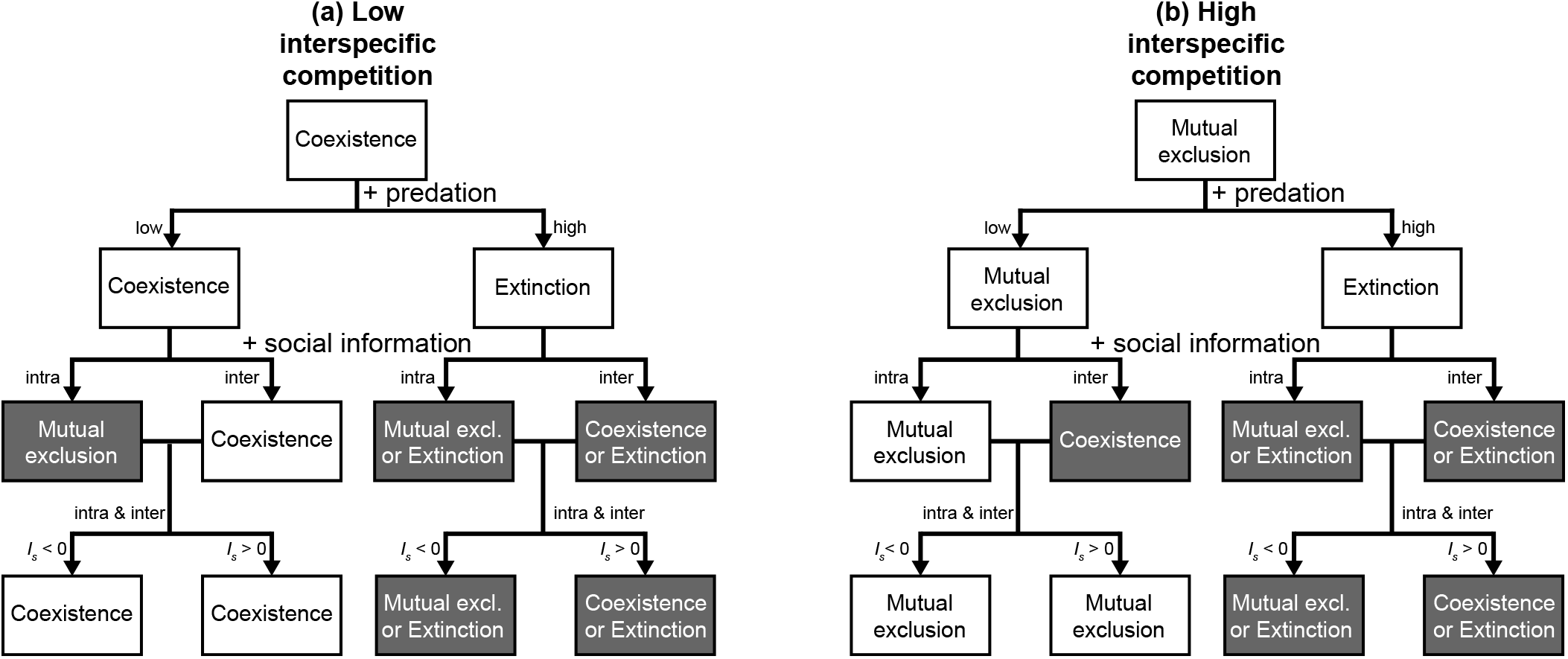
Summary of effects of different types of social information (intraspecific and/or interspecific) on the fate of populations of competing species at equilibrium, under low interspecific competition (a; 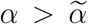) or high interspecific competition (b; 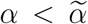), and under low predation (*r* > *p*; left branches) or high predation (*r* < *p*; right branches). Here, when social information is added to the system in either form (3rd row) or both forms (4th row), it is relatively strong (i.e., intra alone: 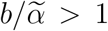 and 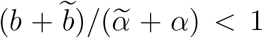, inter alone: 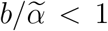 and 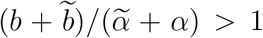, intra & inter: 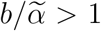 and 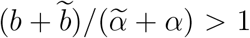, see Appendix S3 for details; Eq. 2; Fig. 2). The inclusion of social information can drive qualitative shifts in the fate of the system (denoted by shaded boxes). Note that such shifts from coexistence to mutual exclusion (a) or vice versa (b) are generally expected at low to intermediate levels of social information (e.g., bottom-left corners of Fig. 6c, d; just beyond the white ‘Extinction’ region in Fig. 6a, b) and become less likely at high levels of both types of social information, unless there is sufficiently high predation or there are strong asymmetries in information types (e.g., see right edges of Fig. 6a, c and top edges of Fig. 6b, d). Qualitative shifts (grey boxes) in the 3rd row follow (from left to right) from Fig. S3-1 aiii, aiv, biii, biv, and qualitative shifts in the 4th row follow from Fig. S3-2 aiv-avii, biv-bvii.

## Appendix S4: Effects of social information on competition in the face of asymmetries

In this Appendix, we use numerical simulations to examine the effects of social information on competitive dynamics between two species when one species is competitively superior, or when one species is preferentially consumed by a shared specialist predator. Note that in both of these cases of asymmetric competing species, the net interaction indices, as presented in the main text, are no longer sufficient to determine the sequence of dynamics the system can exhibit. However, see Appendix S3 for a modification of the inequality used to create *I_s_* (Eq. 5) that accounts for species-specific differences in population growth rate and mortality rate due to predation.

### Superior competitor

To measure the demographic consequences of social information when two competing species are not competitively equivalent (i.e., symmetric), we use Eq. 2 to model species 1 as a superior competitor (*α*_12_ < *α*_21_), whose population, *N*_1_, has an advantage over the population of species 2 (*N*_2_), an inferior competitor, for most initial conditions. When interspecific competition exceeds intraspecific competition (Fig. S1a), species 1 outcompetes species 2 over a greater range of initial conditions (Fig. S1ai), and when intraspecific competition exceeds interspecific competition (Fig. S1b), species 1 and 2 coexist, but *N*_1_ is greater than *N*_2_ over a greater range of initial conditions (Fig. S1bi). These outcomes result from the absence of social information, or from the case when the effects of intraspecific and interspecific social information are equivalent and the same for both species. However, if intraspecific or interspecific social information has a greater effect on one species, this can quantitatively or qualitatively shift competitive outcomes; when social information provides greater benefits to competitively inferior species, these species can persist or reach the larger of the two population sizes over a greater range of initial conditions than competitively superior species (Fig. S1aii, aiv, bii, biv), and when social information provides greater benefits to competitively superior species, these species can exert greater dominance over the inferior competitor, in some cases excluding this species under all initial conditions (Fig. S1aiii, biii).

### Specialist predator

We use Eq. 2 to model the case when the predator shared between the competing populations specializes on (i.e., prefers) one of the two species; in this case, the predator prefers species 1 over species 2 (*p*_1_ > *p*_2_). When interspecific competition exceeds intraspecific competition (Fig. S2a), species 2 can competitively exclude the predator-targeted species 1 over all initial conditions (ai), and when intraspecific competition exceeds interspecific competition (Fig. S2b), species 1 and 2 coexist, with *N*_2_ greater than *N*_1_ (Fig. S2bi). These outcomes result from the absence of social information, or from the case when the effects of intraspecific and interspecific social information are equivalent and the same for both species. However, if intraspecific or interspecific social information has a greater effect on one species, this can quantitatively or qualitatively shift competitive outcomes; when social information provides greater benefits to the prey species that is preferred by the predator (species 1), this species that could otherwise go extinct can persist and reach the larger of the two population sizes over a greater range of initial conditions than the species that is less preferred by the predator: species 2 (Fig. S2aiii, av). When social information provides greater benefits to the species less preferred by the predator (species 2), this species can exert greater dominance over the predator-targeted species (species 1), which can cause or maintain competitive exclusion of this species over all initial conditions (Fig. S2aii, aiv, bii).

**Fig. S1:**
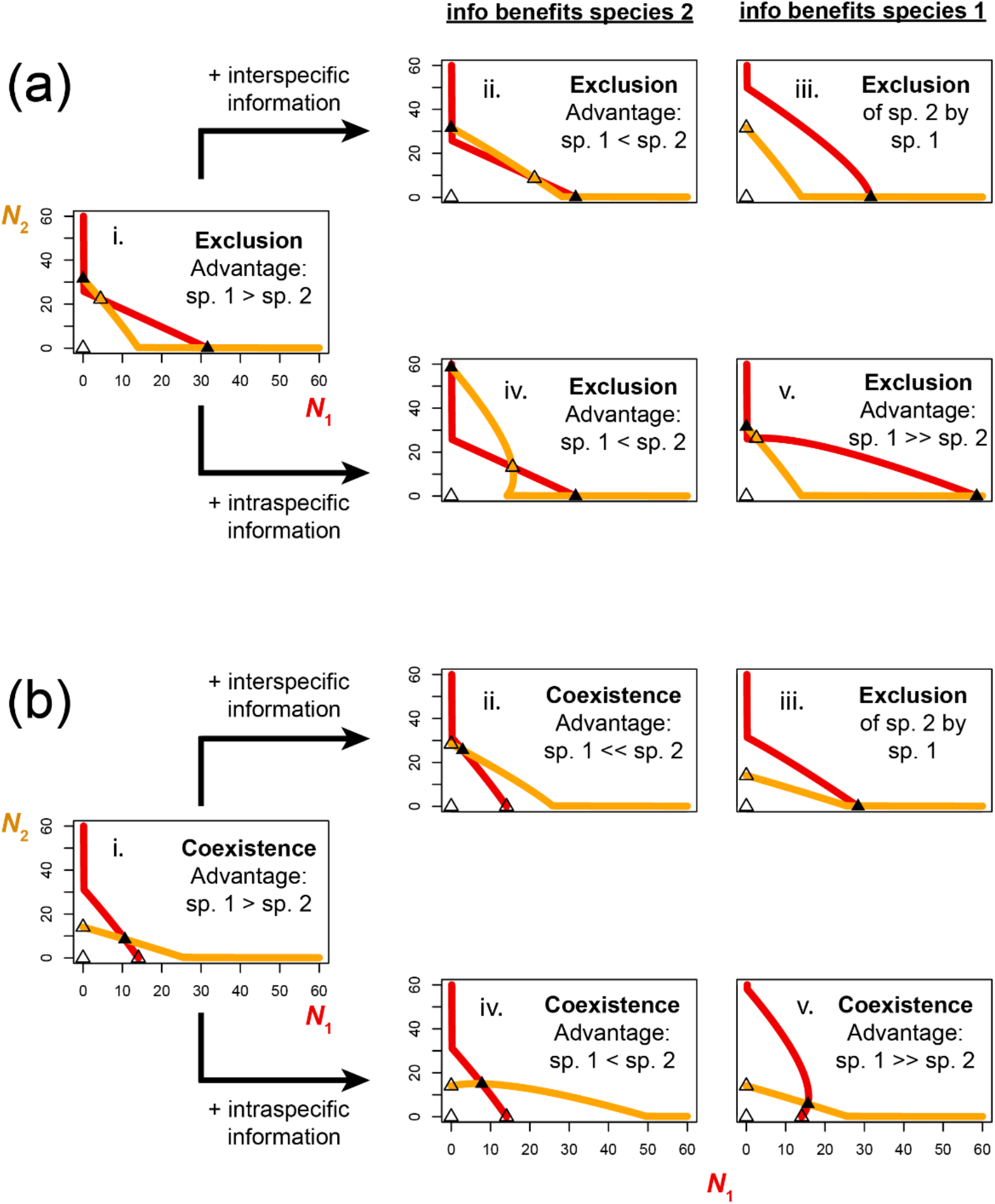
Effects of social information on the outcomes of asymmetric competition. Phase plane plots of nullclines at which the population of each competing species exhibits zero net growth (*N*_1_ nullcline in red, *N*_2_ nullcline in yellow); where nullclines intersect mark equilibrium points (open points denote unstable equilibria; closed points denote stable equilibria). When one species is competitively superior (species 1, in this case), and whether interspecific competition exceeds intraspecific competition (a) or intraspecific competition exceeds interspecific competition (b), asymmetries in effects of social information can determine the outcome of competitive interactions on populations. As in the main text figures, Parameters used: all panels: *r* = 1, *p* = 0.9, (a): α_11_ = α_22_ = 0.01, α_12_ = 0.011, α_21_ = 0.015; (b): α_11_ = α_22_ = 0.015, α_12_ = 0.010; α_21_ = 0.011; (ai, bi): b_11_ = b_22_ = 0.01, b_12_ = b_21_ =0.01; (aii, biv): b_11_ = b_22_ = 0.01, b_12_ = 0.01, b_21_ = 0.02; (aiii, bv): b_11_ = b_22_ = 0.01, b_12_ = 0.02, b_21_ = 0.01; (aiv, bii): b_11_ = 0.01, b_22_ = 0.02, b_12_ = b_21_ = 0.01; (av, biii): b_11_ = 0.02, b_22_ = 0.01, b_12_ = b_21_ = 0.01.

**Fig. S2:**
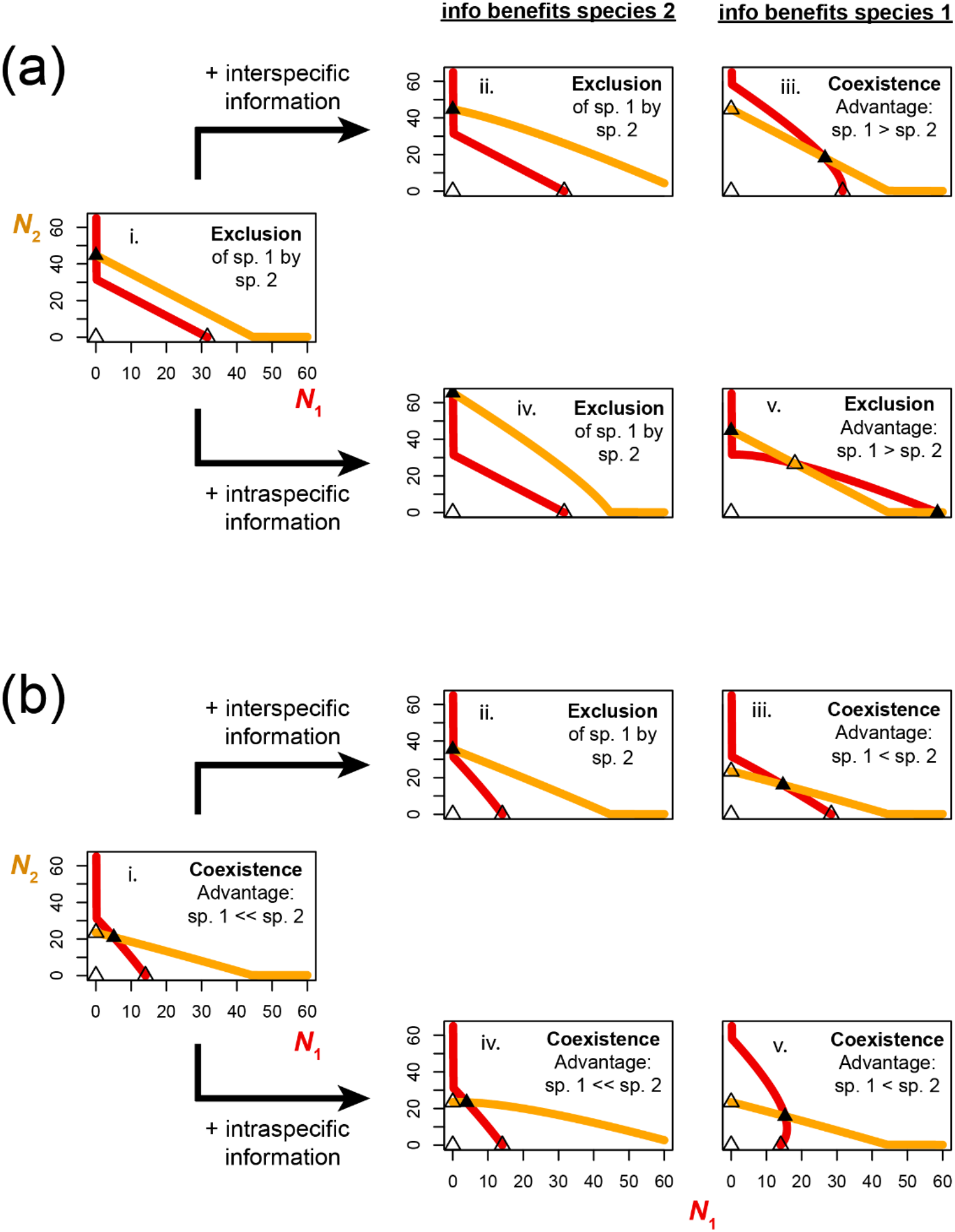
Effects of social information about a specialist predator. Phase plane plots of nullclines at which the population of each competing species exhibits zero net growth (*N*_1_ nullcline in red, *N*_2_ nullcline in yellow); where nullclines intersect mark equilibrium points (open points denote unstable equilibria; closed points denote stable equilibria). When one prey species (species 1, in this case) is preferentially consumed by a predator that is shared by a second, less-preferred species (species 2), and whether interspecific competition exceeds intraspecific competition (a)or intraspecific competition exceeds interspecific competition (b), asymmetries in effects of social information can determine the outcome of competitive interactions on populations. Parameters used: all panels: *r*_1_ = *r*_2_ = 1, *p*_1_ = 0.9, *p*_2_ = 0.8, α_12_ = α_21_ = 0.01; (a): α_11_ = a22 = 0.01; (b): α_11_ = α_22_ = 0.015; (ai, bi): b_11_ = b_22_ = 0.01, b_12_ = b_21_ =0.01; (aii, biv): b_11_ = b_22_ = 0.01, b_12_ = 0.01, b_21_ = 0.02; (aiii, bv): b_11_ = b_22_ = 0.01, b_12_ = 0.02, b_21_ = 0.01; (aiv, bii): b_11_ = 0.01, b_22_ = 0.02, b_12_ = b_21_ = 0.01; (av, biii): b_11_ = 0.02, b_22_ = 0.01, b_12_ = b_21_ = 0.01.

